# Blood RNA Signatures Predict Recent Tuberculosis Exposure in Mice, Macaques and Humans

**DOI:** 10.1101/830794

**Authors:** Russell C. Ault, Colwyn A. Headley, Alexander E. Hare, Bridget J. Carruthers, Asuncion Mejias, Joanne Turner

## Abstract

Tuberculosis (TB) is the leading cause of death due to a single infectious disease. Knowing when a person was infected with *Mycobacterium tuberculosis* (*M.tb*) is critical as recent infection is the strongest clinical risk factor for progression to TB disease in immunocompetent individuals. However, time since *M.tb* infection is challenging to determine in routine clinical practice. To define a biomarker for recent TB exposure, we determined whether gene expression patterns in blood RNA correlated with time since *M.tb* infection or exposure. First, we found RNA signatures that accurately discriminated early and late time periods after experimental infection in mice and cynomolgus macaques. Next, we found a 6-gene blood RNA signature that identified recently exposed individuals in two independent human cohorts, including adult household contacts of TB cases and adolescents who recently acquired *M.tb* infection. Our work supports the need for future longitudinal studies of recent TB contacts to determine whether biomarkers of recent infection can provide prognostic information of TB disease risk in individuals and help map recent transmission in communities.

## Introduction

Tuberculosis (TB) is the leading killer due to a single infectious disease, causing over 1 million deaths per year (*1*). Despite renewed efforts to combat the TB epidemic, the current decline in TB incidence of 1.5% per year has fallen far short of the needed 4-5% annual decline to meet the 2020 goals for the World Health Organization’s (WHO) End TB Strategy (*2*). While approximately ¼ (1.7 billion) of the world’s population has been infected with its causative agent *Mycobacterium tuberculosis* (*M.tb*), only 5 to 10% of infected individuals will develop active TB disease during their lifespan, with the remainder controlling the infection in a state known as latent TB infection (LTBI) (*3, 4*). Recent global workshops have reemphasized targeting transmission of TB as critical to accelerating efforts to reduce the burden of TB disease throughout the world (*5, 6*). Two critical areas for understanding and preventing TB transmission are knowing where and when transmission occurs, and preventing infected individuals from progressing to active TB disease and thereafter transmitting the bacteria via the airborne route (*7, 8*).

Historically, successful control of TB in nations has followed from a reduction in transmission to very low levels (*7, 9*). Studies of close contacts, and in particular household contacts, of active TB cases are a critical tool for identifying new active TB cases from recent transmission and targeting therapy for preventing both subsequent disease and transmission. However, in high incidence countries where most of the burden of disease resides, more than 80% of TB transmission occurs outside of the home (*10, 11*). Genotyping *M.tb* isolates from active TB cases coupled with comparative genomic analysis has permitted population-level identification of hotspots of localized transmission, but these data are mostly available retrospectively and thus do not allow real-time monitoring of TB transmission in a community, particularly in areas of high incidence (*12*). It thus remains unknown whether with current methods TB transmission can be appreciably disrupted in high incidence settings. This is in contrast to low incidence settings where both contact studies and targeting specific higher incidence communities have been effective (*13*).

Recent infection is the single strongest clinical risk factor for developing active TB disease in immunocompetent persons, who comprise the vast majority of LTBI and active TB cases (*14–19*). However, time since exposure or infection is very difficult to ascertain in the clinical setting, and its estimate is often unreliable (*20*). Moreover, there are no known validated biomarkers of recent exposure or infection beyond conversion on a tuberculin skin test (TST) or IFN-γ release assay (IGRA), which requires longitudinal sampling. At the same time, treating all LTBI+ individuals in areas of high TB incidence to prevent the development of active TB is not feasible and would entail unnecessary risk to the vast majority of LTBI+ individuals who will never develop disease. Prospective gene expression-based (RNA) signatures of risk of developing active TB disease have been recently identified for LTBI+ adolescents and adult healthy household contacts (HHCs) (*21, 22*). While the positive predictive value of these RNA signatures of risk of active TB is higher than TST/IGRAs, they are still significantly less than ideal: to prevent one case of active TB, ∼37-64 LTBI+ people not at risk need to be treated (*vs.* ∼85 for TST/IGRA) (*21–23*). It is currently unknown whether these RNA signatures correlate with time since infection. Importantly, their positive predictive value for TB progression and the number needed to treat could be dramatically improved if combined with accurate knowledge of time since infection in the same individual.

Building on this prior work, we assess RNA expression as a potential biomarker of recent exposure or infection with *M.tb*. Using our murine data and recently published studies in cynomolgus macaques and humans (*21, 22, 24, 25*), we show for the first time that RNA expression predicts recent infection/exposure in all three species. Moreover, in both macaques and humans, these RNA signatures of recent infection/exposure are independent of the recently identified signatures of individual prospective TB disease risk. However, in LTBI+ adolescents and adult HHCs, our RNA signature of recent infection was unable to provide prognostic information of TB disease risk, possibly because of its likely duration of only a few months. Our work supports the need for future longitudinal studies of recent TB contacts to identify biomarkers of recent infection that have sufficient duration to provide prognostic information of TB disease risk in individuals and to help map recent transmission in communities.

## Results

### Blood genome-wide RNA expression accurately discriminates early *vs.* late *M.tb* infection time periods in C57BL/6 mice

While several published studies have made genome-scale measurements of the *in vivo* host response to *M.tb* at several time points in mice (*26–28*), none have addressed the question of whether these parameters can predict infection time point. To determine whether it is possible to predict time since *M.tb* infection in mice via a blood RNA signature, we measured genome-wide RNA expression in whole blood in C57BL/6 mice following low dose aerosol *M.tb* infection. Mouse cohorts were sacrificed every month post-infection for 5 months (n = 4 per time point) along with age-matched uninfected mice (n = 1-2 per time point). While *M.tb* colony forming units were not measured, it is well characterized that in this mouse strain lung bacterial burden increases exponentially from the day of *M.*tb infection until the peak of the adaptive immune response in the lungs at 1 month post-infection, thereafter remaining stable for approximately 300 days (*29–31*). Thus, lung CFUs do not predict time since infection in this model after one month post-infection.

Principle component analysis (PCA) of our whole dataset revealed that the blood transcriptional state of *M.tb* infection during the first five months was distinct from that of uninfected mice, with uninfected and infected mice being entirely separable along the 1^st^ principle component (21.8% of data variance; Figure 1A). When we performed PCA on only *M.tb* infected mice, we found that early (30-60 days) and late (90-150 days) time periods were transcriptionally distinct, being separable along the 1^st^ and 2^nd^ principle components (18.8% and 16.5% of data variance, respectively; Figure 1B). Only 1 mouse from the 60 day time point clustered with the late time period along the 2^nd^ principle component.

**Figure 1.**
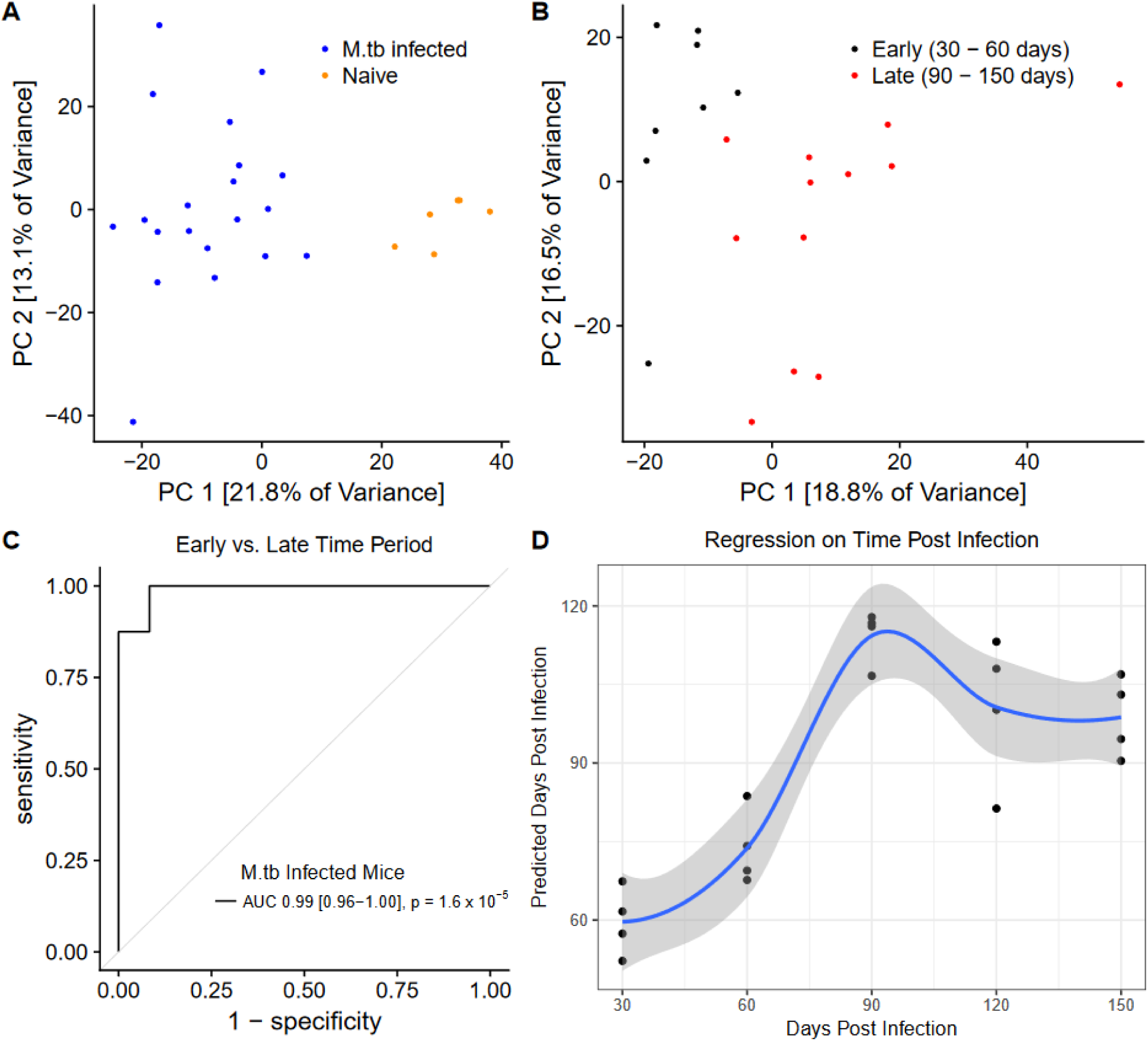
Blood genome-wide RNA expression discriminates early *vs.* late *M.tb* infection time periods in C57BL/6 mice. Principle component analysis of genome-wide RNA expression measured via microarray in (A) all mice (n = 6 uninfected mice, n = 20 *M.tb* infected mice) stratified by infection status and (B) only *M.tb* infected mice stratified by time period post-infection. (C) ROC curve for out-of-bag performance of Random Forest Classifier predicting time period post-infection (1-2 months *vs.* 3-5 months; *P* from Wilcoxon test, 95% confidence interval shown). (D) Random Forest Regression out-of-bag predictions of monthly time point post-infection. Fit curve calculated via the Loess method with 95% CI shown.

To find a predictive RNA signature of time since *M.tb* infection, we used the Random Forest Classifier algorithm, without hyperparameter tuning due to low sample size, to predict early (30-60 days) *vs.* late (90-150 days) infection time period. Using out-of-bag predictions (approximately 3-fold cross-validation) to obtain an unbiased estimate of predictive performance, we found that we could predict early *vs.* late infection time period with 0.99 area under the curve (AUC) (95% CI: 0.96 – 1.00, *P* = 1.6 x 10^-5^; 87.5% sensitivity, 91.7% specificity for early infection; Figure 1C). To assess whether each month post-infection could be predicted accurately, we performed Random Forest Regression with 3-fold cross-validation and confirmed that days 30 and 60 were predicted to be earlier time points than days 90-150 (Figure 1D). Days 90-150 were not resolved. Although low group size precludes confident quantification of the degree to which days 30 and 60 can be separated, 3 out of 4 mice within both early time points were predicted in the correct order. Probes used in these models as well as their feature importance for the regression model are shown in Table S1. Taken together, these data indicate that we can broadly discriminate early and late *M.tb* infection in this cohort of C57BL/6 mice based on the whole blood transcriptomic response, and it may even be possible to discriminate between the first two months of infection.

### Blood RNA signature discriminates early *vs.* late *M.tb* infection time periods in cynomolgus macaques

While inbred mice are a suitable model for studying molecular components of the immune response to *M.tb*, they do not replicate the variable clinical outcomes of *M.tb* infection in humans. Cynomolgus macaques, an outbred non-human primate model for TB, do exhibit heterogeneity in clinical outcomes, with approximately half of macaques progressing to symptomatic active TB disease that can be verified radiologically and bacteriologically within the first 6 months of infection, and the remainder controlling the infection in a latent state (*32, 33*). The lung pathology of *M.tb* infection in cynomolgus macaques also better replicates several features of human lung pathology than mice (*33*).

To determine whether our findings in the murine model translated to the more human-like cynomolgus macaque model of *M.tb* infection, we mined publically available data from a longitudinal study of *M.tb* infection in macaques (*24*). In that study, cynomolgus macaques were infected with a low dose of *M.tb* in the lung, and their blood was sampled at 11 time points post-infection and 2 time points pre-infection for genome-wide RNA expression analysis. Importantly, while the study’s authors provided a broad, unsupervised analysis of their data according to time periods of infection, they did not assess our hypothesis that blood genome-wide RNA expression predicts time period or time point post-infection (*24*). To test our hypothesis and allow comparison with our mouse data and recently available human data, we restricted our analysis to 8 time points from 20 days through 180 days (6 months) post-infection. To permit comparison of different computational models and allow a final unbiased estimate of predictive performance, we randomly divided the 38 macaques from this study into a training set and a test set, keeping the ratio of macaques with latent and active TB balanced in both groups (Figure S1).

Using 9-fold cross-validation on the training set, we found that Regularized Logistic Regression, a linear method, was not inferior to several nonlinear classification methods in predicting early (20-56 days) *vs.* late (90-180 days) infection time period (Figure S2). We thus chose Regularized Logistic Regression to find a predictive RNA signature of time since *M.tb* infection in cynomolgus macaques. We found that this model predicted early (20-56 days) *vs.* late (90-180 days) infection time period with an AUC of 0.78 in the training set (95% CI: 0.72-0.85, *P* = 5.6 x 10^-13^; 9-fold cross-validation; Figure 2A), and an AUC of 0.81 in the test set (95% CI: 0.71-0.91, *P* = 1.6 x 10^-7^; Figure 2A).

**Figure 2.**
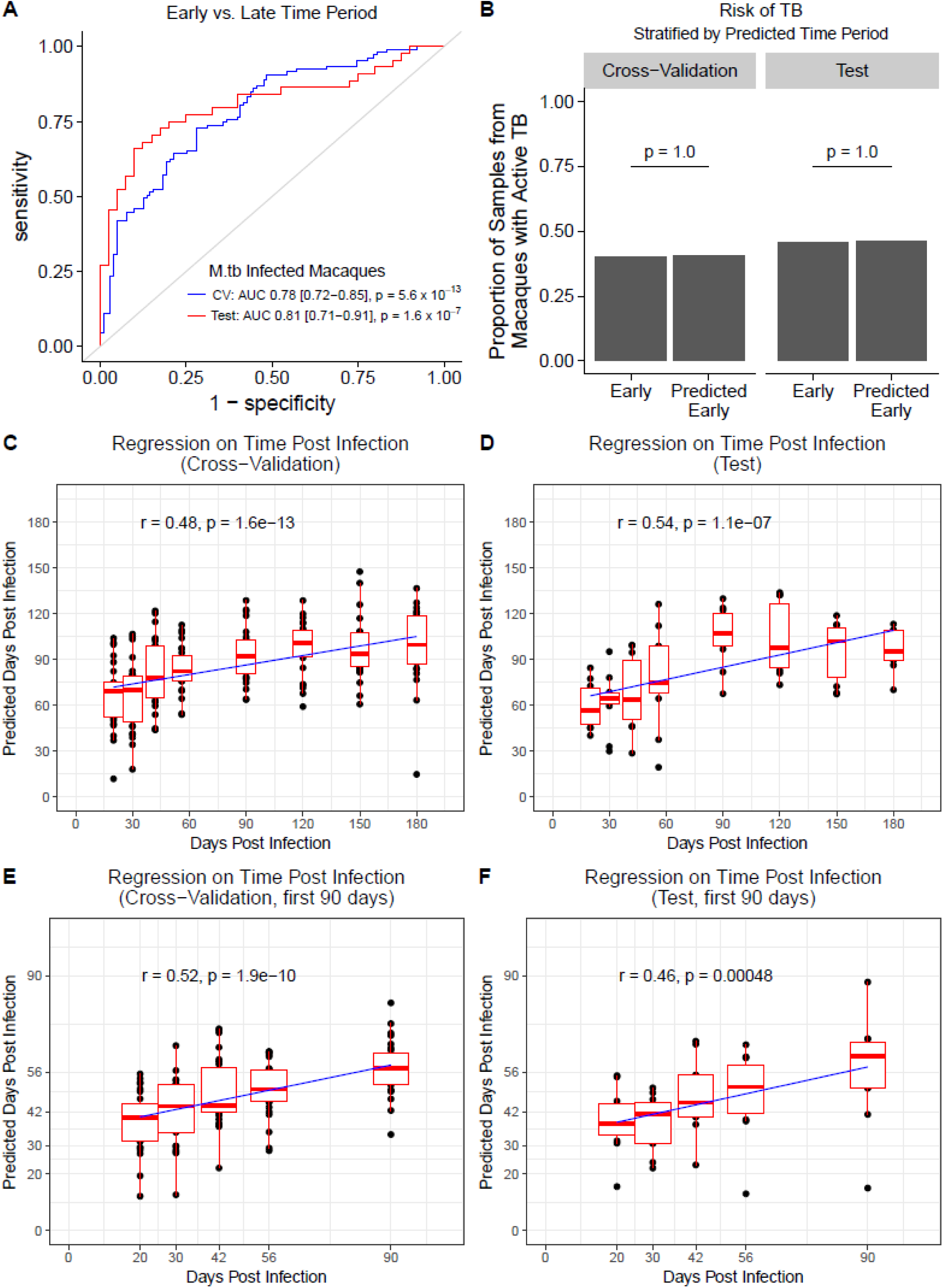
Blood RNA signature discriminates early *vs.* late *M.tb* infection time periods in cynomolgus macaques. (A) ROC curves for Regularized Logistic Regression prediction of time period post-infection (20-56 days *vs.* 90-180 days) from RNA expression in cynomolgus macaques on 9-fold cross-validation in the training set (blue curve; n = 107 early time period samples, n = 103 late time period samples) and final model prediction on test set (red curve; n = 44 early time period, n = 40 late time period) (*P* from Wilcoxon test). (B) Comparison between early (20-56 days) (n = 107 train, n = 44 test) *vs.* predicted early (n = 104 train, n = 50 test) time period samples in proportion of samples from macaques that develop active TB (*P* from Fischer’s Exact test). Regularized Linear Regression predictions of time point post-infection for (C) 9-fold cross-validation in the training set (n = 210) and for (D) final model prediction on the test set (n = 84). (E-F) Predictions from models trained and evaluated only on samples from the first 90 days post-infection (n = 134 train, n = 55 test). Boxplots represent medians with interquartile ranges for the predictions at each time point (best fit line shown, *P* from Pearson test).

Importantly, our model was trained and tested on macaques irrespective of their present or future TB disease status. If our model partially predicted disease status rather than only time period post-infection, the proportion of samples from macaques with active disease would differ between predicted and actual early time period samples. However, we found that there was no change in the proportion of samples from macaques with active disease in the predicted early time periods relative to the actual early time periods, in both the training and test sets (*P* = 1.0, *P* = 1.0 respectively; Figure 2B). This was also true focusing on late time period predictions (*P =* 1.0 training, *P =* 1.0 test; data not shown).

Next, to assess whether each month post-infection could be predicted in cynomolgus macaques, we performed Regularized Linear Regression with 9-fold cross-validation on the training set and confirmed that days 20-56 were predicted as earlier time points than days 90-180, in both the training set and in the test set (Figure 2C-D). As in our murine model analysis days 90-180 were not resolved. Quantitatively, the median absolute error (MAE) of the model was 38.5 days (Pearson’s *r* = 0.48, *P =* 1.6×10^-13^) on the training set and 35.7 days (*r* = 0.54, *P =* 1.1×10^-7^) on the test set. Probes selected and used by the final trained regression model to predict in the test set are shown in Table S2. To assess whether we could predict specific time point of infection within the first 3 months, as suggested by our murine data, we trained a model on only time points from 20-90 days (Figure 2E-F). The MAE of this model was 15.8 days on the training set and 14.3 days on the test set (*r* = 0.52, *P* = 1.9×10^-10^ and *r* = 0.46, *P* = 4.7×10^-4^, respectively).

Taken together, these data indicate that we can broadly discriminate early and late *M.tb* infection in this cohort of cynomolgus macaques based on the whole blood transcriptomic response, and that we can moderately discriminate between the first two months of infection. These predictions do not depend on disease status, and the accuracy of the predictions is quantitatively lower in cynomolgus macaques than in C57BL/6 mice, as reflected by the AUC analyses.

### Blood RNA expression of 250 genes predicts time since active TB exposure in humans

To determine whether our findings in mice and cynomolgus macaques could translate to humans, several points are important to consider. While a recent study in the United States and Canada showed that recent contacts of active TB cases are at highest risk of TB disease in the first 1-3 months after the diagnosis of the TB index case, studies in other countries and other time periods show that the highest risk is in the first 1-2 years, with most cases accruing more than 2-3 months after a documented exposure (*14, 17, 34–37*). A recent vaccine study in rhesus macaques showed that BCG-induced immunity to *M.tb* delays IGRA conversion in a repeated limiting-dose *M.tb* challenge model (*38*). Therefore, the natural course of infection or reinfection and RNA correlates of recent infection in humans could be the same or delayed relative to our analysis in models of infection in *M.tb*-naïve animals. This could depend on local transmission burden and the likelihood of pre-existing immunity to *M.tb*, whether from BCG or a prior *M.tb* exposure (*39*).

Whereas the day of infection is known in animal models, the precise time of exposure resulting in infection is difficult to determine in humans, even in careful clinical studies. One surrogate for time of infection in humans is time of IGRA or TST conversion in people who were known to be IGRA/TST negative previously. This would synchronize a human study cohort to the time of an initial systemic T cell response to *M.tb*. To test this hypothesis we accessed public data from South African adolescents who acquired latent *M.tb* infection during longitudinal blood sampling every 6 months (*25*). We found that Regularized Logistic Regression was unable to predict the first time point of known IGRA conversion from 6 months post-first known IGRA conversion (0.54 AUC, 95% CI: 0.27-0.82, *P* = 0.64 on test set; Figure 3A). Notably, the biological event of actual IGRA conversion in this cohort could have occurred anytime between the first time point of IGRA positivity and the preceding 6 months. Given our findings in mice and macaques that the RNA signature of time since *M.tb* infection occurs within a brief window of 2-3 months, we interpret these findings to mean that sampling blood every 6 months in humans is unlikely to constitute a cohort where actual time of IGRA conversion is synchronized sufficiently to discover an RNA signature of time since IGRA conversion.

**Figure 3.**
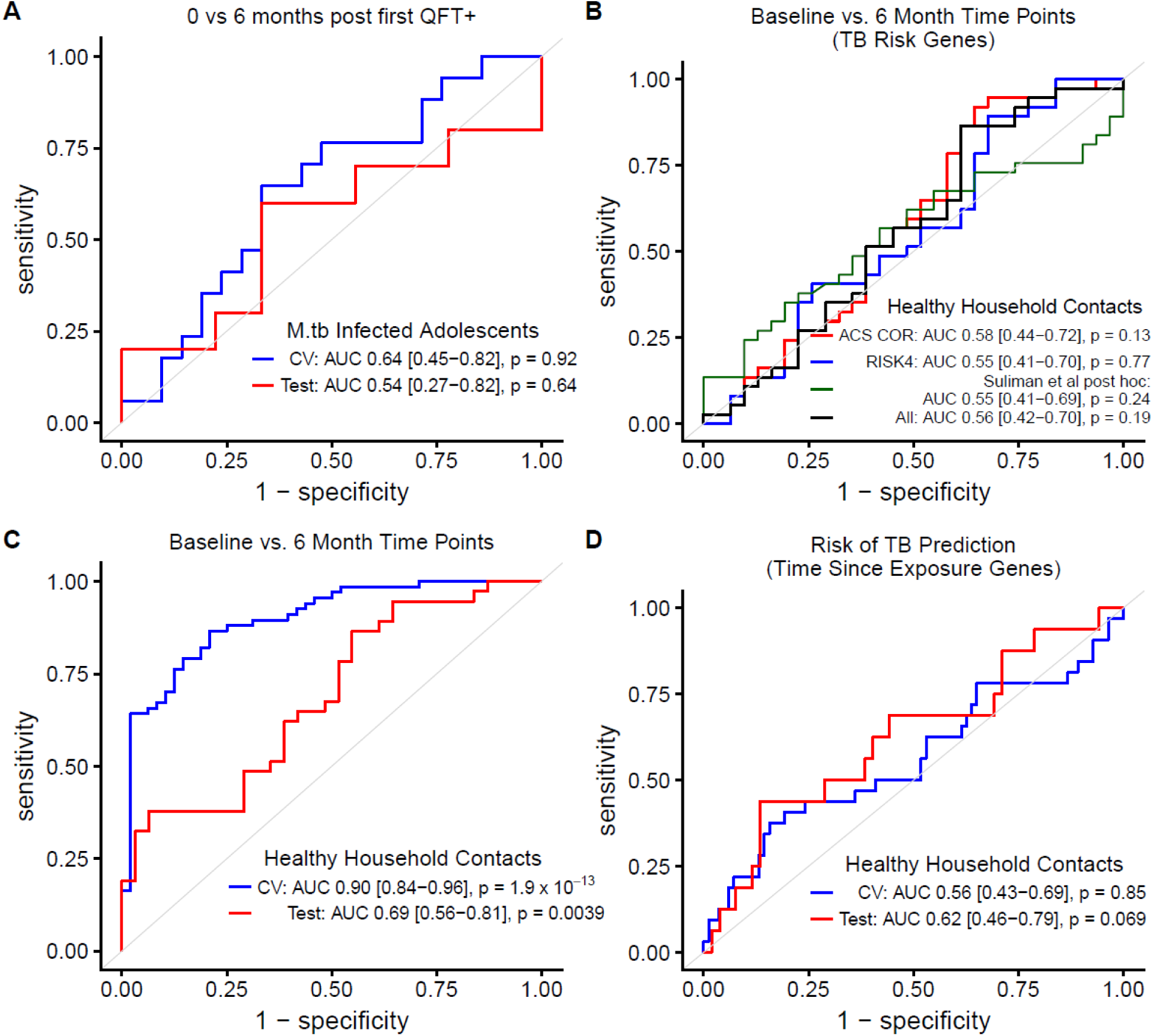
Blood RNA expression of 250 genes predicts time since active TB exposure in humans. (A) ROC curves for prediction of time since first known IGRA+ (0 *vs.* 6 months) in South African adolescents who acquire *M.tb* infection for 10-fold cross-validation in the training set (blue curve; n = 17 0 month samples, n = 21 6 month samples) and final model prediction on the test set (red curve; n = 10 0 month, n = 9 6 month) using Regularized Logistic Regression. (B) ROC curves for Regularized Logistic Regression prediction of time since active TB exposure (baseline *vs.* 6 months post-enrollment) in GC6-74 Gambia and Ethiopia test set (n = 37 baseline samples, n = 31 6 months samples) using expression of genes from published signatures that predict prospective risk of active TB. (C) ROC curves for Regularized Logistic Regression prediction of time since active TB exposure for 10-fold cross-validation on the Gambia and Ethiopia training set (blue curve; n = 67 baseline, n = 48 6 months) and for final model prediction (contains 250 genes) on the Gambia and Ethiopia test set (red curve; n = 37 baseline, n = 31 6 months). (D) ROC curves for prediction of prospective risk of TB for 10-fold cross-validation on the Gambia and Ethiopia training set (blue curve; n = 67 baseline, n = 48 6 months) and for final model prediction on the test set (red curve; n = 37 baseline, n = 31 6 months) using the 250-gene set that predicted time since active TB exposure. *P* values for all ROC curves are from Wilcoxon test, and 95% confidence intervals are shown.

Another study design that could identify RNA correlates of recent infection in humans is a household contact study wherein healthy contacts of active TB cases are enrolled within a certain time from the date of diagnosis of the active TB case and sampled longitudinally. Important limitations of this design that could reduce the power to detect RNA correlates of recent infection are that the precise time of infection is not known and individuals who are IGRA+ at enrollment may have been infected either from the present exposure or in the more distant past. Cognizant of these limitations, we accessed publically available data from the Grand Challenges 6-74 (GC6-74) study of healthy household contacts (HHCs) of patients with active pulmonary TB (*22*). HHCs in this cohort were enrolled within 2 months of the diagnosis of the active TB index case and had blood samples drawn at baseline, 6 months and/or 18 months post-enrollment (*22*). Because our mouse and macaque analysis suggests that blood transcriptional changes are most prominent in an early 3 month window post-infection, we focused our first analysis on the baseline and 6 month time points. This included data from Gambian and Ethiopian cohorts but excluded data from the South African cohort because 6 month time points were not available for South Africa (*22*). We used the same training/test split as the authors in the Gambian cohort but randomly split the Ethiopian cohort 50/50 between our training and test sets. Importantly, with this training/test split and our data pre-processing, we could predict risk of TB with 0.72 AUC (95% CI: 0.60-0.83, *P* = 1.6 x 10^-4^; data not shown) in the training set by 10-fold cross-validation and 0.70 AUC (95% CI: 0.53-0.88, *P* = 0.0071; data not shown) in the test set using Regularized Logistic Regression. From the GC6-74 and the Adolescent Cohort Study (ACS) we used the RISK4 genes (*BLK*, *CD1C*, *GAS6* and *SEPT4*), the post-hoc selected *C1QC*, *TRAV27*, *ANKRD22*, *OSBPL10* genes and the 16 correlate of risk (COR) predictive genes together for this analysis (*21, 22*). When we used these same genes to train a model to predict time since TB exposure, we obtained no predictive performance, whether the model was trained with these gene sets separately or together (*P* > 0.05 for all test set predictions; Figure 3B). This suggests that genes selected for optimal prediction of prospective TB risk do not change across these two time points post-exposure.

To find a predictive RNA signature of time since TB exposure in these data, and as the study authors performed for TB risk prediction, we used the Wilcoxon test on the training set to select transcripts that differed in expression between baseline and 6 month time points (*22*). Using Regularized Logistic Regression we found that these genes predicted baseline *vs.* 6 month time points with 0.90 AUC (95% CI: 0.84-0.96, *P* = 1.9×10^-13^; 10-fold cross-validation; Figure 3C) in the training set and 0.69 AUC (95% CI: 0.56-0.81, *P* = 0.0039; Figure 3C) in the test set. We further used the final genes selected by the model (250 genes, Table S3) on the training set to train a model to predict risk of TB. As expected, these genes exhibited no direct predictive performance for risk of TB on the training or test sets (*P* = 0.85, *P* = 0.07, respectively; Figure 3D). In summary, our findings with the household contact study design in humans parallel the results in macaques in that we can predict broad time period post-exposure via the whole blood transcriptomic response. Moreover, this transcriptomic signature of time period post-exposure to an active TB case is independent of the transcriptomic signature of risk of TB recently identified in the GC6-74 and ACS studies (*21, 22*).

### Time since TB exposure in humans is associated with alteration in CD4+ T cell proportion and immune activation pathways

Cell-type deconvolution algorithms have recently been used with genome-wide RNA expression data to help identify changes in immune cell proportions in the blood that are associated with TB disease, prospective TB disease risk and treatment success (*25, 40*). To identify immune cell populations that are associated with time since TB exposure in the GC6-74 study, we used the leukocyte expression signature matrix ‘immunoStates’ and linear regression to infer leukocyte proportions for each subject’s sample (*40*). We found that the proportion of CD4+ α/β T cells was increased at 6 months *vs.* baseline time point in the Gambian and Ethiopian cohorts (*P* = 0.0079; linear mixed model; Figure 4A), but was not significantly changed at 18 months (*P* = 0.20 *vs.* baseline; linear mixed model, included South African cohort; Figure 4B). We saw no significant differences in NK cell proportion over time in the Gambian and Ethiopian cohorts (Figure 4C-D). Likewise, no other cell types estimated by the ‘immunoStates’ signature matrix showed significant differences over time in these cohorts (*P* > 0.05, linear mixed model, data not shown). This result with CD4+ α/β T cells and NK cells is consistent with the conclusion that the RNA signature of time since TB exposure is independent from the RNA signature of prospective TB risk, since both T cells and NK cells are known to decrease in circulation in active TB disease (*25, 41*).

**Figure 4.**
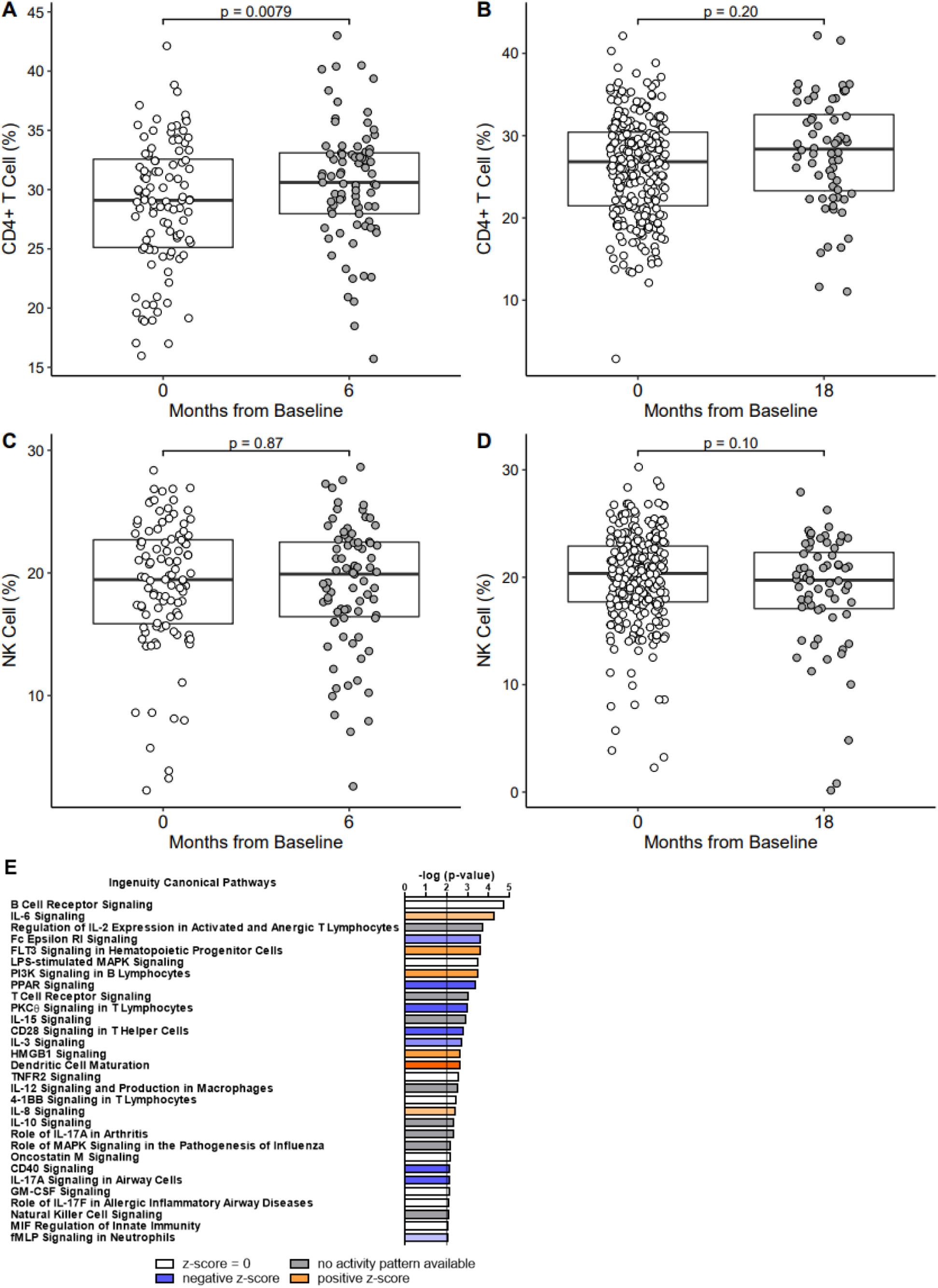
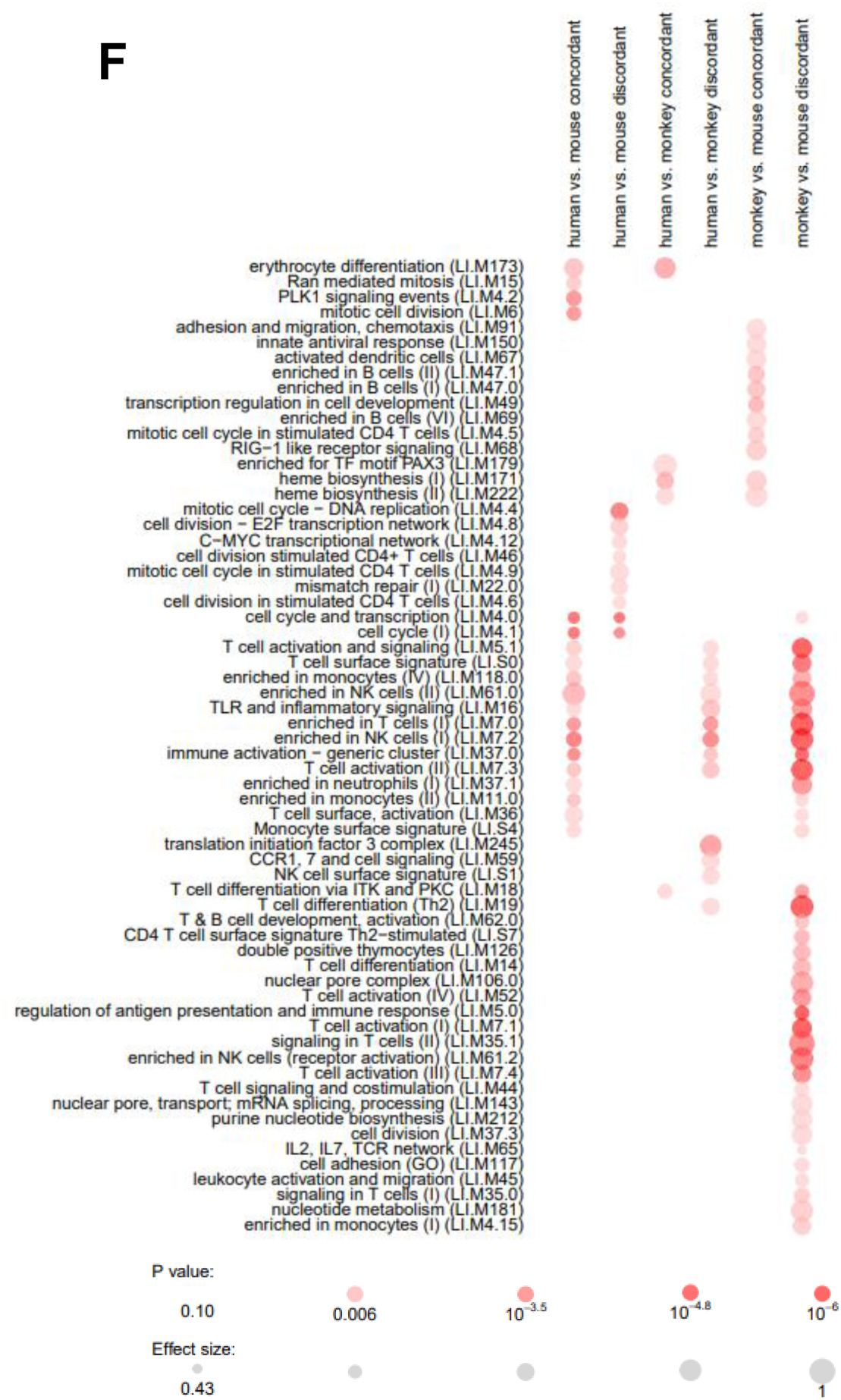
Time since TB exposure in humans is associated with alteration in CD4+ T cell proportion and immune activation pathways. Changes in CD4+ T cell percentages (A,B) and NK cell percentages (C, D) in GC6-74 healthy household contacts cohort at baseline (n = 104 in A,C; n = 272 in B, D), 6 month (A, C; n = 79) and 18 month (B, D; n = 64) time points after active TB exposure were determined by cell-type deconvolution (*P* from linear mixed model). Boxplots represent medians with interquartile ranges. (E) Top immunity related enriched canonical pathways in the 250-gene RNA signature of time since exposure to active TB index case (6 months *vs.* baseline) by IPA (*P* from Fisher’s Exact test). (F) Enriched transcriptional modules that are concordantly or discordantly regulated during recent *M.tb* exposure or infection between mice, macaques or humans by disco analysis (*P* from CERNO statistical test).

Our RNA signature of baseline *vs.* 6 month time points post-exposure included 250 genes selected by Regularized Logistic Regression (Table S3). We utilized Ingenuity Pathway Analysis (IPA) to identify pathways associated with these genes. The majority of enriched canonical pathways (-log(p-value)>2) were associated with immune cell signaling, including B cells (B cell receptor and PI3K signaling), T cells (T cell receptor, PKCθ, regulation of IL-2 expression, 4-1BB and CD28 signaling), cytokines (IL-6, IL-15, IL-12, TNF, IL-8, IL-10 and IL-17A), innate immune cells (dendritic cell maturation and LPS-stimulated MAPK signaling) and humoral immunity (Fc Epsilon RI Signaling) (Figure 4E, Table S4). Other enriched canonical pathways were related to cellular injury and toxicity (apoptosis), metabolism, nervous system signaling, PPAR signaling, cell cycle regulation and intracellular & second messenger signaling (Table S4).

Considering the overall direction of change in the immune pathways between 6 month *vs.* baseline time points, the upregulation of several pro-inflammatory signaling pathways (IL-6, IL-8, FLT3 signaling, PI3K signaling in B Lymphocytes and Dendritic Cell maturation) and decrease in anti-inflammatory signaling (PPAR signaling) suggests that an increase in peripheral blood immune activation occurs at the 6 month time point after exposure (Figure 4E).

To compare transcriptional modules altered in humans to those altered in mice and macaques, we used the recently defined disco score to identify concordantly and discordantly altered modules between these species (*42*). Several modules related to T cells, NK cells and monocytes were enriched (adjusted *P* < 0.05) in each pairwise comparison between two species (human *vs.* mouse, human *vs.* monkey, and monkey *vs.* mouse) (Figure 4F). Several B cell-related modules were uniquely concordantly regulated between macaques and mice (Figure 4F).

### Application of reduced 6-gene expression signature of time since active TB exposure to adolescent *M.tb* infection acquisition cohort confirms its identification of recent infection in humans

Implementation of our newly discovered RNA signature of time since active TB exposure using qRT-PCR would require a more parsimonious gene set than the 250 genes heretofore described. To find a reduced gene signature we ran a forward search using the MetaIntegrator R package (*43*). This method identified 6 genes, *RP11-552F3.12, PYURF, TRIM7, TUBGCP4, ZNF608* and *BEAN1*, that recapitulated the performance of the 250 gene signature on baseline *vs.* 6 month time point discrimination with 0.86 AUC (95% CI: 0.80-0.93, *P* = 1.7×10^-11^; Figure 5A) in our GC6-74 training set and 0.68 AUC (95% CI: 0.55-0.81, *P* = 0.0055; Figure 5A) in the test set. Independent validation of this signature requires a cohort wherein recent *M.tb* infection is documented and time points are available to test whether the signature allows discrimination between recent and more remote infection. While the cohort of South African adolescents who acquired latent *M.tb* infection did not permit discovery of an RNA signature of recent *M.tb* infection, we reasoned that the whole cohort would be powered for validation of our signature discovered in the GC6-74 household contact study design (*25*). Three genes, *TRIM7*, *ZNF608*, *TUBGCP4*, from our 6-gene signature were represented by detected probes in the microarray used in this study. These 3 genes discriminated the first time point of known IGRA conversion from all pre-conversion time points (6 months and 12 months prior to known conversion) with 0.72 AUC (95% CI: 0.58-0.87, *P* = 0.0030; Figure 5B). These 3 genes likewise discriminated the first time point of known IGRA conversion from all sampled time points (6, 12 months prior to conversion and 6, 12 months after known conversion) with 0.68 AUC (95% CI: 0.56-0.81, *P* = 0.0039; Figure 5B). Figure S3 shows the trajectory of the 3 gene score over time, being highest at the first time point of known IGRA conversion.

**Figure 5.**
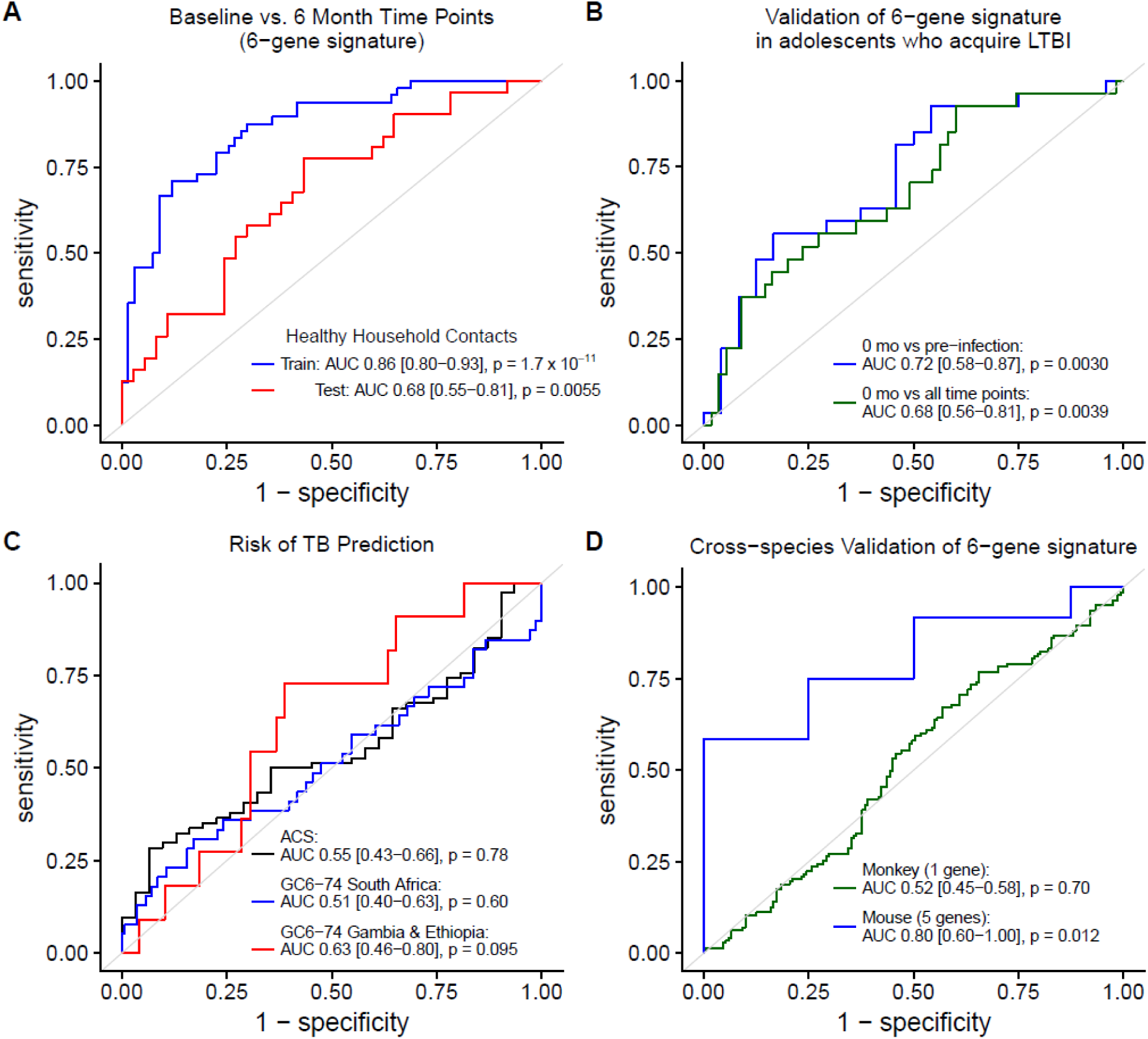
Application of reduced 6-gene signature of time since active TB exposure to adolescent *M.tb* infection acquisition cohort confirms its identification of recent infection in humans. (A) ROC curves for 6-gene score prediction of time since active TB exposure in the Gambia and Ethiopia training set (blue curve; n = 67 baseline samples, n = 48 6 months samples) and for the Gambia and Ethiopia test set (red curve; n = 37 baseline, n = 31 6 months). (B) ROC curves for discrimination between time of first known IGRA+ and all pre-conversion time points (blue curve; n = 27 0 month, n = 24 pre-conversion) and between time of first known IGRA+ and all other time points (green curve; n = 27 0 month, n = 24 pre-conversion and n = 31 6 or 12 months after known conversion) in South African adolescents who acquire *M.tb* infection using 3-gene score from genes detected in microarray data. (C) ROC curves for prediction of prospective risk of TB using highest 6-gene score observed per individual in the ACS cohort (n = 74 nonprogressors, n = 31 progressors), GC6-74 Gambia and Ethiopia test set (n = 49 nonprogressors, n = 11 progressors) and GC6-74 South Africa cohort (n = 141 nonprogressors, n = 39 progressors). (D) ROC curves for discrimination of early and late time periods post-infection in mice (blue curve; n = 8 early mice, n = 12 late mice) and macaques (green curve; n = 151 early samples, n = 143 late samples) using genes from the 6-gene signature that were detected in the respective microarrays. *P* values for all ROC curves are from Wilcoxon test, and 95% confidence intervals are shown.

Given that time since active TB exposure is the single strongest clinical risk factor for developing TB disease in immunocompetent persons, the finding that time since exposure and risk of TB, as predicted by the blood transcriptomic response, are independent in the GC6-74 study of healthy household contacts suggests that these signatures could be combined to possibly better predict risk of TB when the time of exposure is unknown (*17, 19, 20*). While the GC6-74 study was not powered for this particular secondary analysis, we assessed whether the highest 6-gene score during longitudinal sampling allowed discrimination of subjects who did or did not progress to active TB disease during study follow-up. The highest 6-gene score did not discriminate progressors from non-progressors in the test set, whether the subjects were from South Africa (AUC 0.51, 95% CI: 0.40-0.63, *P* = 0.60; Figure 5C) or from Gambia or Ethiopia (AUC 0.63, 95% CI: 0.46-0.80, *P* = 0.095; Figure 5C). The same result was observed in the ACS cohort of IGRA+ adolescents with unknown exposure history (AUC 0.55, 95% CI: 0.43-0.66, *P* = 0.78; Figure 5C).

We additionally tested whether the 6-gene signature discriminated early from late time periods post-infection as defined in our analysis of animal models of *M.tb* infection. The 5 genes represented by detected probes of homologous genes in the mouse microarray data discriminated the early (30-60 days) *vs.* late (90-150 days) infection time period with 0.80 AUC (95% CI: 0.60 – 1.00, *P* = 0.012; Figure 5D). Only 1 gene was represented by a detected probe in the macaque microarray data, and it alone did not discriminate the early *vs.* late infection time period (0.52 AUC, 95% CI: 0.45 – 0.58, *P* = 0.70; Figure 5D). Of note, this study utilized a human microarray platform for the macaque samples, which may have contributed to reduced measurability of the macaque homologues to these human genes (*24*).

## Discussion

Early clinical studies in the pre-antibiotic era in the relatively isolated Faroe Islands shed light on the clinical features of primary infection with *M.tb* in humans, which often include fever, elevated erythrocyte sedimentation rate, X-ray abnormalities and, less often, erythema nodosum (*34, 44*). With time of exposure to an active TB case pinpointed within a two week period, and sometimes to a single day, Poulsen determined that these clinical features accompany TST conversion within 6 weeks of exposure (*14, 34, 44*). While these clinical features of initial *M.tb* infection are transient and not specific to *M.tb* infection, a method to determine that a person is currently in the first 1-2 years post initial infection would have great prognostic value for near-future TB disease and could allow real-time geospatial mapping of recent TB transmission in communities (*14, 15, 17, 34–37*). In our proof of concept analysis, we sought to determine whether it is possible to develop an RNA-based blood test to detect recent exposure or infection with *M.tb*. For TB disease risk prediction, we hypothesized that such a test could complement the recently developed RNA signatures of TB disease risk that are based on detecting incipient tuberculosis, which is the asymptomatic phase of early TB disease during which pathology progresses gradually before full-blown clinical TB (*21, 22, 45, 46*).

Using our mouse data and published macaque data, we have demonstrated a highly accurate RNA signature of recent infection with *M.tb* (1-2 *vs.* 3-5/6 months post-infection). Using the GC6-74 cohort of HHCs of patients with active pulmonary TB, we discovered 250-gene and 6-gene human RNA signatures of recent exposure (0-2 *vs.* 6-8 months post-diagnosis of index case) that validated within a held-out test set (*22*). Using an independent cohort of adolescents who acquired *M.tb* infection during 6-month longitudinal sampling, we demonstrated that the 6-gene signature could discriminate the first known time point of IGRA conversion from pre-conversion time points and from 6-12 months later with modest accuracy (0.68 AUC) (*25*).

However, this 6-gene signature was unable to provide prognostic information of TB risk in the GC6-74 cohort or the ACS cohort. The incomplete time point sampling of most individuals, and 6-month sampling likely reduced the power to find an association between our 6-gene signature score and TB risk in these two studies. Nevertheless, we believe the sampling constraints and target populations of these studies, adults who are HHCs and adolescents with LTBI of unknown exposure history, both in highly endemic areas, are mostly in line with what may be feasible for applying transcriptional signatures of TB risk for targeted treatment to reduce TB incidence (*46*). Given that early blood transcriptional changes occurred within a short 3 month window in our mouse and macaque analyses, and the human data analyzed are not inconsistent with this brief timeline, we believe that blood RNA signatures for recent *M.tb* infection are too brief in duration to yield a useful biomarker to improve prediction of TB risk for targeted preventive therapy.

A recent genetic study of early progression to TB disease (within 18 months) demonstrated that the genetic architecture of early progression and later reactivation disease are different (*47*). Because the vast majority of TB disease burden can be accounted for epidemiologically by recent infection (past 1-2 years), we hypothesize that, on average, the genetic and environmental factors influencing progression of disease have resolved by 2 years post initial infection in humans (*3, 14*). Therefore, we hypothesize that a biological correlate of recent infection that has the longest duration during that time when the outcome of early disease progression has not been resolved would have the highest chance of being useful as a complement to tests for incipient TB in predicting TB risk. Indeed, our estimated cell type and pathway analyses suggest that both cellular and molecular signatures of immune activation associated with recent exposure and could be interrogated by other modalities such as epigenetics. Immune cell differences between recently acquired and remotely acquired infection have been reported by others in single cohorts without longitudinal sampling (*20, 48*). The high enrichment of B cell signaling in our signature is interesting, and a recent case control study in a single cohort showed that several IgG and IgA antibodies to *M.tb* antigens strongly discriminated (AUC > 0.90) active TB contacts who converted on TST from non-converters both at first known conversion and 3 months prior (*49*). Using dense sampling (> 1/month) for 3 years in an individual, DNA methylation was recently shown to have more prolonged dynamics in human blood in response to chronic disease states than RNA expression, and thus represents an epigenetic modality to be considered (*50*).

Our analyses and these considerations suggest that sampling IGRA-, untreated HHCs every month (or more frequently) for one to two years, starting as soon as possible after the diagnosis of their respective index case and determining IGRA conversion events, would allow for the discovery of biosignatures of recent *M.tb* infection that could be useful for helping predict TB disease risk. Follow-up in such a cohort for TB progression would allow better assessment of how signatures of recent infection and signatures of incipient TB could be combined to improve TB risk prediction. The addition of chest X-rays with deep machine learning analysis could be useful to discover heretofore unknown, specific radiogenomic features of recent infection or incipient TB (*51, 52*). After IGRA conversion, staggered sampling at different times could reduce the study’s burden on individual subjects and allow more precise estimation of the duration of any biomarker. Most follow-up in such a study would have to be performed on those who refuse preventive treatment, as treatment would need to be offered because recent infection is precisely documented. Another potential benefit of such a study is that validated biomarkers that associate strongly with TST/IGRA conversion but precede conversion, such as currently unvalidated IgG and IgA markers, could be used to identify *M.tb* infection before TST/IGRA conversion and thus reduce the burden of follow-up of recent contacts in TB control programs and potentially help reduce LTBI treatment time (*49*).

If deployed in population screening efforts, a test for recent *M.tb* infection could also allow real-time geospatial mapping of recent TB transmission in communities. This could greatly help the application of current control methods to reduce TB transmission and disease in high incidence settings. While it is possible that our current 6-gene signature of recent *M.tb* infection could be evaluated in the future for this purpose, we think it would be more prudent to first find biomarkers of recent *M.tb* infection that have a longer duration and are useful for individual TB risk prediction. However, biomarkers of varying duration could be jointly useful for the application of mapping recent transmission.

Our results in mice, macaques and humans, together with recent literature, suggest that future longitudinal studies of HHCs may be successful at identifying more accurate biomarkers of time since *M.tb* infection in humans. Our study represents one of only a handful of studies since Poulsen’s early work showing that there are biological events in the early human response to *M.tb* infection that can be reproducibly measured (*34*). Future biomarker studies may enable the study of early events of infection in humans both routinely and ethically and permit the identification of immunological or other biological events that determine whether an exposed person will develop TB disease or control the infection (*5, 53–55*). This could greatly aid vaccine development for TB as no correlates of protection for TB are yet known (*56*). We also expect that more accurate biomarkers of time since *M.tb* infection will be excellent tools to help better understand the human phenotypes of IGRA reversion and persistent resistance to IGRA conversion (*54, 57*).

Our current analysis has some limitations. Because most transmission occurs outside the household contact setting, many individuals in the GC6-74 study were TST+ at enrollment (∼51.4% in Ethiopia, ∼36.3% in The Gambia), and follow-up TST in this study were incomplete, it is highly likely that many, and possibly the majority, of contacts in this study were not infected or re-infected from their index TB case (*10, 11, 58*). However, the 6-gene RNA signature discovered in this cohort validated in adolescents where recent *M.tb* infection was documented via IGRA conversion in 100% of study participants (*25*). Finally, our current analysis excluded HIV co-infection.

## Methods

### Study Design

The objective of this study was to identify blood RNA correlates of time since *M.tb* infection or exposure. We first infected mice with *M.tb* via the aerosol route and measured genome-wide RNA expression at pre-specified time points. Unsupervised analysis revealed potential discrimination between mice sacrificed at early time points (1-2 months) *vs.* late time points (3-5 months). Cross-validation without hyperparameter tuning identified an unbiased RNA signature that accurately predicted early *vs.* late time period post-infection. We then retrospectively mined data from a prospective *M.tb* infected cynomolgus macaque cohort and a prospective healthy household contact human cohort to identify RNA signatures that predicted these same time periods post-infection. The human RNA signature was validated in an independent cohort, adolescents who were recently infected with *M.tb* during longitudinal sampling.

### Mice

Specific pathogen-free, 6-12 week old, female C57BL/6 wild-type mice (The Jackson Laboratory, Bar Harbor, ME) were maintained in ventilated cages inside a biosafety level 3 (BSL3) facility and provided with sterile food and water *ad libitum*. All protocols were approved by The Ohio State University’s Institutional Laboratory Animal Care and Use Committee.

### Mouse aerosol infection and blood collection

*M.tb* Erdman (ATCC no. 35801) was obtained from the American Type Culture Collection. Stocks were grown according to published methods (*59*). Mice were infected with *M.tb* Erdman using an inhalation exposure system (Glas-Col) calibrated to deliver 50 to 100 CFUs to the lungs of each mouse, as previously described (*59, 60*). At specific time points post-*M.tb* infection, infected and age-matched uninfected mice were sacrificed and blood collected (400 μL) from the heart into 1.2 mL Tempus reagent and stored at −80°C. No formal randomization was employed for choosing cages of mice to be sacrificed at each time point. For the *M.tb* infected mice, sample size per time point was determined by using the number we routinely use for well-powered molecular and immunological studies in inbred mice. No blinding was performed for the mouse study.

### RNA processing and microarray hybridization

Whole blood RNA was processed, quantified using a NanoDrop 1000™ spectrophotometer (NanoDrop Technologies) and RNA integrity (RIN) determined by a 2100 Bioanalyzer (Agilent). Samples with RIN ≥ 6.5 were submitted for hybridization onto Illumina mouse WG 6-V2 beadchips and scanned on an Illumina Beadstation system. Microarray data will become available in the Gene Expression Omnibus (GEO) database upon publication.

### Microarray data pre-processing

For our murine data, Illumina BeadStudio/GenomeStudio software was used to subtract background and scale average signal intensity for each sample to the global median average intensity across all samples. Probes with a detection *P* value ≤ 0.01 in at least 10% of mice were filtered for analysis. Thereafter R scripts were used to quantile normalize the data, set all values <10 to 10 and log_2_ transform the data. Probes were filtered by two-fold change in expression from the median in at least 10% of samples. For the macaque data (GSE84152), microarray data pre-processing was performed as previously described (*24*). The data from the human adolescent cohort of IGRA converters (GSE116014) was pre-processed identically as the macaque data, except that data were quantile normalized and no batch correction was performed. When these adolescent data were used to validate the 6-gene signature, the data were downloaded at the gene-level using the R MetaIntegrator package, before additional pre-processing (*43*).

### RNA-seq data pre-processing

Human data from the Grand Challenges 6-74 (GC6-74) cohort were downloaded at the gene count level from GEO (GSE94438). Genes with read count ≤ 5 in 50% of samples were excluded. Data were quantile normalized and log_2_ transformed. To facilitate comparisons with a common RNA-seq alignment pipeline, gene counts were obtained from the ARCHS^4^ resource when comparing data from the Adolescent Cohort Study (GSE79362) and GC6-74 cohorts using the 6-gene signature (*61*). These data were otherwise processed identically.

### Machine learning predictions

For predicting time since infection in mice, we used the Random Forest algorithm in R with default parameter values (*62*). Out-of-bag predictions were used to estimate model accuracy, which corresponds approximately to 3-fold cross-validation.

To predict time since infection in macaques, we randomly partitioned the macaques into training (70%) and test (30%) sets. We compared several different machine learning algorithms using the R caret package (*63*). These included: Random Forest (R ranger package (*64*)), Gradient Boosted Machines (R gbm package (*65*)), Support Vector Machines using Polynomial (R kernlab package (*66*)) or RBF kernels (R kernlab package (*66*)) and Regularized Logistic Regression (R glmnet package (*67*)). 9-fold cross validation was used in the training set to optimize model hyperparameters and assess predictive performance, with all samples related to individual macaques being partitioned into the same held-out fold to ensure unbiased cross-validation. The caret package implementation did not permit 10-fold cross validation for this dataset, as in humans, but the results should be equivalent. Only Regularized Logistic Regression was used for predictions in the test set and Regularized Linear Regression for predicting each time point post-infection after Regularized Logistic Regression was shown to be superior in predicting time period post-infection.

To predict time since TB exposure, time since IGRA conversion or prospective risk of TB in humans, we used 10-fold cross validation on the training set (either GC6-74 or Adolescent IGRA converter cohort), with each subject’s samples partitioned into the same held-out fold, to optimize Regularized Logistic Regression model hyperparameters before predicting on the test set. Prior to performing this procedure for time since TB exposure on the GC6-74 training set, we performed feature selection on genes by a Wilcoxon test (*P* < 0.05). Where longitudinal data were available for individual macaques or persons, each time point was considered as an independent sample.

### Forward search to discover parsimonious 6-gene signature

A forward search was performed in the GC6-74 Gambia and Ethiopia training set on genes selected by a Wilcoxon test (*P* < 0.05) using the R MetaIntegrator package as previously described (*43, 68*). The stopping threshold for increase in AUC with the addition of each gene was varied until a signature comprising less than 10 genes and including both upregulated and downregulated genes at 6 months post-enrollment (*vs.* baseline) was obtained. The final signature’s score is calculated on normalized log_2_ expression values as a difference between upregulated and downregulated genes: (*RP11-552F3.12 + PYURF + TRIM7 + TUBGCP4*) – (*ZNF608* + *BEAN1*). When applying this score to microarray data, multiple detected probes that mapped to these genes, using the R biomaRt package, were averaged (*69*). Genes without corresponding detected probes were omitted from the calculation.

### Cell type deconvolution, pathway and transcriptional module analysis

Cell type proportions in blood were estimated from RNA-seq data as previously described using the R MetaIntegrator package (*25, 40, 43*). Gene-level expression for this deconvolution was obtained from the ARCHS^4^ resource (*61*). For pathway analysis, the 250 genes comprising the signature of time since exposure to an active TB case (6 months *vs.* baseline) were analyzed using canonical pathway analysis with QIAGEN’s Ingenuity^®^ Pathway Analysis platform (IPA^®^, QIAGEN Redwood City, www.qiagen.com/ingenuity). To compare transcriptional modules that were concordantly or discordantly regulated between mice, macaques and humans at early and late post-exposure time points, we used the R disco and tmod packages with transcriptional modules from Li et al. (*70–72*). Genes used in this analysis included all detected probes (mice and macaques) and genes (humans). Differential expression and ortholog assignment were performed as previously described (*42*). The 6-month time point in the GC6-74 cohort was taken as the early time point in humans based on the results in Figures 5 and S3, as this time point had the highest 6-gene score (data not shown).

### Statistical analysis

All statistical analyses were performed in R (version 3.4.3). Prediction performance was evaluated using receiver operator characteristic (ROC) curves. Statistical significance of the area under the curve (AUC) was assessed using the one-sided Wilcoxon test via the R Verification package (*73*). ROC graphs and confidence intervals were obtained via the R pROC package (*74*). Pearson test was used for correlation analysis. Fischer’s exact test (two-sided) was used to determine statistical significance of comparisons between proportions in evaluating the independence of the time since infection signatures from risk of TB disease in macaques. We used linear mixed models to assess the significance of cell type proportion changes with time since TB exposure via the R lme4 package (*75*). Subject and site were included as random effects and time since exposure and site as fixed effects. These two-sided *P* values were obtained via the Satterthwaite approximation. The IPA canonical pathway *P* values were calculated by a one-sided Fisher’s Exact Test, with *P <* 0.01 considered as significant. The transcriptional module *P* values were calculated using the CERNO statistical test, with *P* < 0.05 considered as significant after Benjamini-Hochberg correction (*42*). For all other statistical tests, *P* < 0.05 was considered as significant.

### Data availability

Mouse microarray data will be made available in the Gene Expression Omnibus database upon publication. Published data used in this study are available in the Gene Expression Omnibus database under accession numbers GSE79362, GSE84152, GSE94438 and GSE116014.

### Code availability

Source code for all analyses is publicly available in a GitHub repository: https://github.com/remi10001/TB.

## Supplementary Materials

Figure S1. Training and test set partition for cohort of cynomolgus macaques.

Figure S2. Comparison of different machine algorithms to predict time period of *M.tb* infection in cynomolgus macaques.

Figure S3. Trajectory of 3-gene signature for recent *M.tb* infection before and after IGRA conversion in adolescents who acquire *M.tb* infection.

Table S1. Feature importance for all probes used in predicting time since *M.tb* infection in mice (regression of 1-5 months post-infection).

Table S2. Probes comprising 50-probe RNA signature of time since *M.tb* infection in cynomolgous macaques (regression of 1-6 months post-infection).

Table S3. Genes comprising 250-gene RNA signature of time since exposure to active TB index case (6 months *vs.* baseline).

Table S4. Top significantly enriched canonical pathways in 250-gene RNA signature of time since exposure to active TB index case (6 months *vs.* baseline) by IPA.

## Acknowledgments

We thank the personnel of the Ohio State University Animal BSL3 laboratory for assistance, including J. Cyktor for technical help with the mouse experiments. We acknowledge the efforts of the Baylor Institute for Immunology Research Genomics core for assistance with sample preparation and microarray processing. We thank E. Kautto and Q. Hassan for consultation with the mouse data analysis. We thank L. Schlesinger and L. Barreiro for critical review of the manuscript.

## Author contributions

R.C.A. conceived the idea, designed the study and data analysis, analyzed the data and wrote the manuscript. C.A.H. performed the IPA analysis. A.E.H. contributed to the macaque data analysis. B.J.C. performed the mouse experiment. B.J.C. and A.M. prepared samples for RNA microarray analysis. J.T. oversaw the study and data analysis. R.C.A., C.A.H. and J.T. revised the manuscript. All authors commented on the manuscript.

## Corresponding author

Correspondence to Joanne Turner.

## Figures

**Figure S1.**
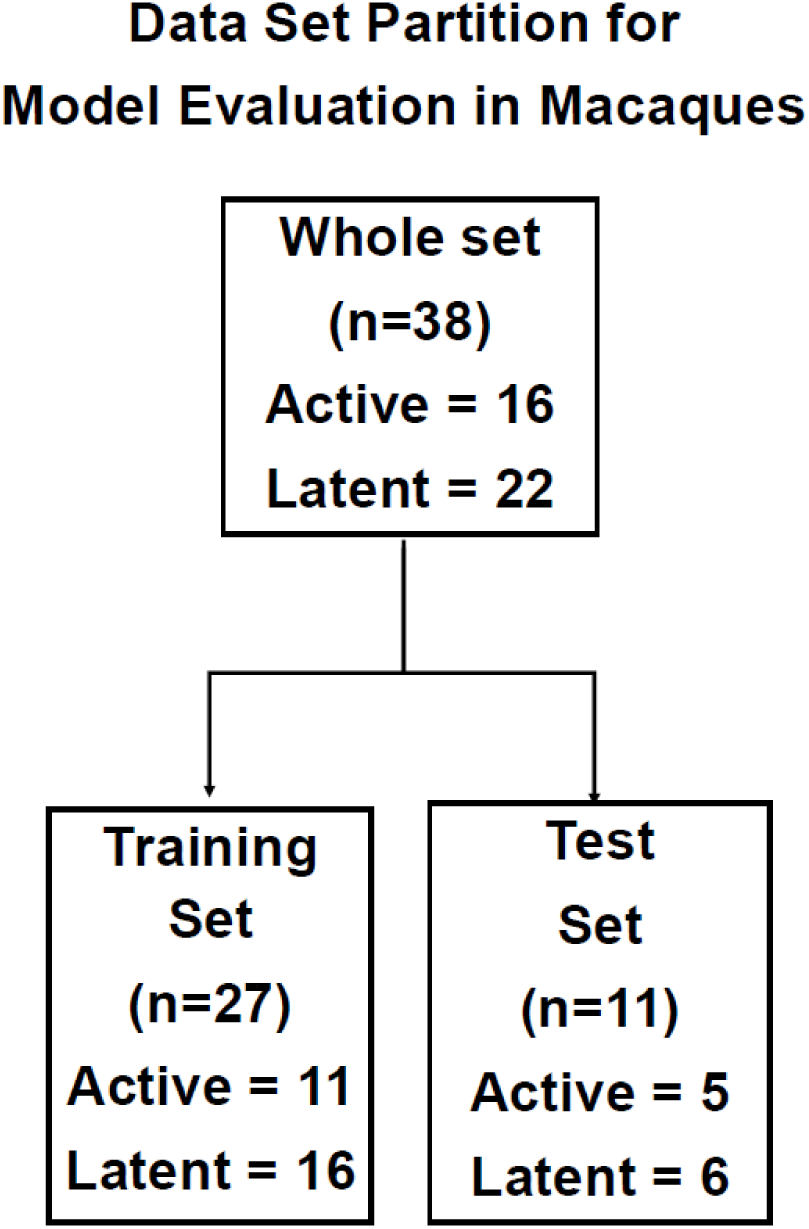
Training and test set partition for cohort of cynomolgus macaques. Active = Developed active TB during the 6 months of study follow-up. Latent = did not develop active TB in this study. n corresponds to individual macaques, each of which underwent longitudinal sampling.

**Figure S2.**
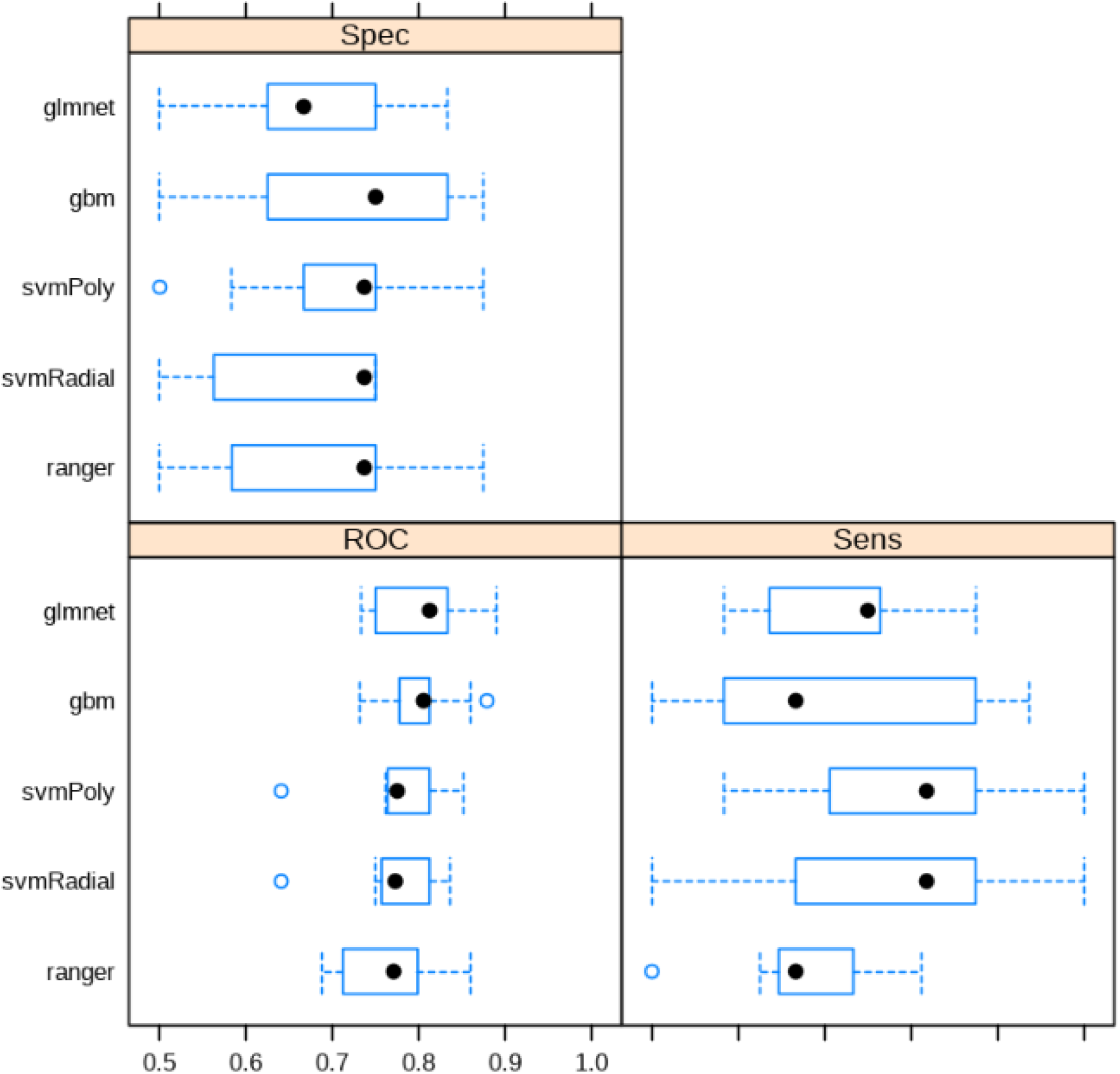
Comparison of different machine algorithms to predict time period of *M.tb* infection in cynomolgus macaques. Random hyperparameter search and 9-fold cross-validation of macaques were used on the training set to evaluate models to predict time period of infection from microarray data. Median (point), interquartile ranges (boxes), and ranges (whiskers) are shown for predictions on each independent fold for the best performing model for each algorithm. glmnet = Regularized Logistic Regression, gbm = Gradient Boosted Machines, svmPoly = Support Vector Machines with Polynomial kernel, svmRadial = Support Vector Machines with RBF kernel, ranger = Random Forest. Sens=Sensitivity, Spec=Specificity, ROC= Area under the curve.

**Figure S3.**
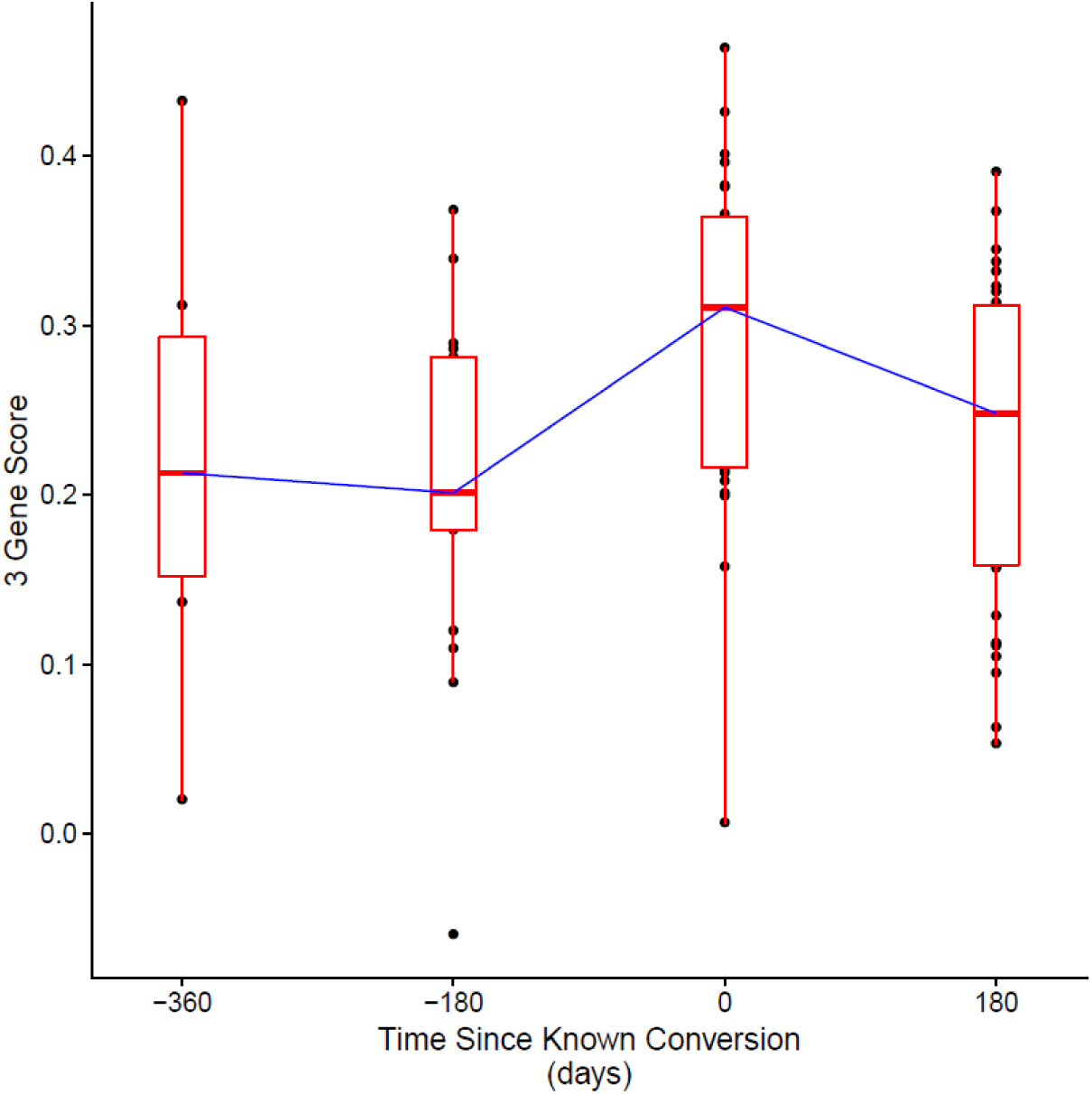
Trajectory of 3-gene signature for recent *M.tb* infection before and after IGRA conversion in adolescents who acquire *M.tb* infection. One sample (score = 0.47) from 360 days after known conversion is omitted but was included in analyses of Figure 5B. Boxplots represent medians with interquartile ranges, and the blue line connects medians. n = 7-360 days, n = 17-180 days, n = 27 0 days, n = 30 180 days.

## References

1. World Health Organization, Global Tuberculosis Report 2018 (World Health Organization, Geneva, Switzerland; http://www.who.int/tb/publications/global_report/en/), p. (WHO, Geneva, 2018).

2. World Health Organization, Global strategy and targets for tuberculosis prevention, care and control after 2015 (World Health Organization, Geneva, Switzerland; http://www.who.int/tb/post2015_strategy/en/), p. (WHO, Geneva, 2014).

3. R. M. G. J. Houben, P. J. Dodd, The Global Burden of Latent Tuberculosis Infection: A Re-estimation Using Mathematical Modelling, PLOS Med. 13, e1002152 (2016).

4. E. Vynnycky, P. E. M. Fine, Lifetime Risks, Incubation Period, and Serial Interval of Tuberculosis, Am. J. Epidemiol. 152, 247–263 (2000).

5. S. Keshavjee, F. Amanullah, A. Cattamanchi, R. Chaisson, K. M. Dobos, G. J. Fox, H. E. Gendelman, R. Gordon, A. Hesseling, H. Le Van, B. Kampmann, B. Kana, G. Khuller, D. M. Lewinsohn, D. A. Lewinsohn, P. L. Lin, L. L. Lu, G. Maartens, A. Owen, M. Protopopova, J. Rengarajan, E. Rubin, P. Salgame, E. Schurr, J. A. Seddon, S. Swindells, D. M. Tobin, Z. Udwadia, G. Walzl, S. Srinivasan, R. Rustomjee, P. Nahid, Moving toward Tuberculosis Elimination. Critical Issues for Research in Diagnostics and Therapeutics for Tuberculosis Infection, Am. J. Respir. Crit. Care Med. 199, 564–571 (2018).

6. N. S. Shah, P. Kim, B. D. Kana, R. Rustomjee, Getting to Zero New Tuberculosis Infections: Insights From the National Institutes of Health/US Centers for Disease Control and Prevention/Bill & Melinda Gates Foundation Workshop on Research Needs for Halting Tuberculosis Transmission, J. Infect. Dis. 216, S627–S628 (2017).

7. G. Churchyard, P. Kim, N. S. Shah, R. Rustomjee, N. Gandhi, B. Mathema, D. Dowdy, A. Kasmar, V. Cardenas, What We Know About Tuberculosis Transmission: An Overview, J. Infect. Dis. 216, S629–S635 (2017).

8. D. W. Dowdy, A. D. Grant, K. Dheda, E. Nardell, K. Fielding, D. A. J. Moore, Designing and Evaluating Interventions to Halt the Transmission of Tuberculosis, J. Infect. Dis. 216, S654–S661 (2017).

9. H. G. Wiker, T. Mustafa, G. A. Bjune, M. Harboe, Evidence for waning of latency in a cohort study of tuberculosis, BMC Infect. Dis. 10, 37–46 (2010).

10. S. Verver, R. M. Warren, Z. Munch, M. Richardson, G. D. van der Spuy, M. W. Borgdorff, M. A. Behr, N. Beyers, P. D. van Helden, Proportion of tuberculosis transmission that takes place in households in a high-incidence area, The Lancet 363, 212–214 (2004).

11. T. A. Yates, P. Y. Khan, G. M. Knight, J. G. Taylor, T. D. McHugh, M. Lipman, R. G. White, T. Cohen, F. G. Cobelens, R. Wood, D. A. J. Moore, I. Abubakar, The transmission of Mycobacterium tuberculosis in high burden settings, Lancet Infect. Dis. 16, 227–238 (2016).

12. J. L. Zelner, M. B. Murray, M. C. Becerra, J. Galea, L. Lecca, R. Calderon, R. Yataco, C. Contreras, Z. Zhang, J. Manjourides, B. T. Grenfell, T. Cohen, Identifying Hotspots of Multidrug-Resistant Tuberculosis Transmission Using Spatial and Molecular Genetic Data, J. Infect. Dis. 213, 287–294 (2016).

13. J. P. Cegielski, D. E. Griffith, P. K. McGaha, M. Wolfgang, C. B. Robinson, P. A. Clark, W. L. Hassell, V. A. Robison, K. P. Walker, C. Wallace, Eliminating Tuberculosis One Neighborhood at a Time, Am. J. Public Health 103, 1292–1300 (2013).

14. M. A. Behr, P. H. Edelstein, L. Ramakrishnan, Revisiting the timetable of tuberculosis, BMJ 362, k2738 (2018).

15. G. J. Fox, S. E. Barry, W. J. Britton, G. B. Marks, Contact investigation for tuberculosis: a systematic review and meta-analysis, Eur. Respir. J. 41, 140–156 (2013).

16. P. Kasaie, J. R. Andrews, W. D. Kelton, D. W. Dowdy, Timing of Tuberculosis Transmission and the Impact of Household Contact Tracing. An Agent-based Simulation Model, Am. J. Respir. Crit. Care Med. 189, 845–852 (2014).

17. M. R. Reichler, A. Khan, T. R. Sterling, H. Zhao, J. Moran, J. McAuley, P. Bessler, B. Mangura, I. Bakhtawar, C. LeDoux, J. McAuley, J. Beison, M. Fitzgerald, M. Naus, M. Nakajima, N. Schluger, Y. Hirsch-Moverman, J. Moran, H. Blumberg, J. Tapia, L. Singha, E. Hershfeld, B. Roche, B. Mangura, A. Sevilla, T. Sterling, T. Chavez-Lindell, F. Maruri, S. Dorman, W. Cronin, E. Munk, A. Khan, Y. Yuan, B. Chen, F. Yan, Y. Shen, H. Zhao, H. Zhang, P. Bessler, M. Fagley, M. Reichler, M. Reichler, T. Sterling, J. Tapia, C. Hirsch, C. Luo, Risk and Timing of Tuberculosis Among Close Contacts of Persons with Infectious Tuberculosis, J. Infect. Dis. 218, 1000–1008 (2018).

18. R. Sloot, M. F. Schim van der Loeff, P. M. Kouw, M. W. Borgdorff, Risk of Tuberculosis after Recent Exposure. A 10-Year Follow-up Study of Contacts in Amsterdam, Am. J. Respir. Crit. Care Med. 190, 1044–1052 (2014).

19. I. Sutherland, Recent studies in the epidemiology of tuberculosis, based on the risk of being infected with tubercle bacilli, Adv. Tuberc. Res. Fortschritte Tuberkuloseforschung Progres Explor. Tuberc. 19, 1–63 (1976).

20. A. Halliday, H. Whitworth, S. H. Kottoor, U. Niazi, S. Menzies, H. Kunst, S. Bremang, A. Badhan, P. Beverley, O. M. Kon, A. Lalvani, Stratification of Latent Mycobacterium tuberculosis Infection by Cellular Immune Profiling, J. Infect. Dis. 215, 1480–1487 (2017).

21. D. E. Zak, A. Penn-Nicholson, T. J. Scriba, E. Thompson, S. Suliman, L. M. Amon, H. Mahomed, M. Erasmus, W. Whatney, G. D. Hussey, D. Abrahams, F. Kafaar, T. Hawkridge, S. Verver, E. J. Hughes, M. Ota, J. Sutherland, R. Howe, H. M. Dockrell, W. H. Boom, B. Thiel, T. H. M. Ottenhoff, H. Mayanja-Kizza, A. C. Crampin, K. Downing, M. Hatherill, J. Valvo, S. Shankar, S. K. Parida, S. H. E. Kaufmann, G. Walzl, A. Aderem, W. A. Hanekom, A blood RNA signature for tuberculosis disease risk: a prospective cohort study, The Lancet 387, 2312–2322 (2016).

22. S. Suliman, E. G. Thompson, J. Sutherland, J. Weiner, M. O. C. Ota, S. Shankar, A. Penn-Nicholson, B. Thiel, M. Erasmus, J. Maertzdorf, F. J. Duffy, P. C. Hill, E. J. Hughes, K. Stanley, K. Downing, M. L. Fisher, J. Valvo, S. K. Parida, G. van der Spuy, G. Tromp, I. M. O. Adetifa, S. Donkor, R. Howe, H. Mayanja-Kizza, W. H. Boom, H. M. Dockrell, T. H. M. Ottenhoff, M. Hatherill, A. Aderem, W. A. Hanekom, T. J. Scriba, S. H. E. Kaufmann, D. E. Zak, G. Walzl, G. Walzl, G. F. Black, G. van der Spuy, K. Stanley, M. Kriel, N. Du Plessis, N. Nene, T. Roberts, L. Kleynhans, A. Gutschmidt, B. Smith, N. Nene, A. G. Loxton, N. N. Chegou, G. Tromp, D. Tabb, T. H. M. Ottenhoff, M. R. Klein, M. C. Haks, K. L. M. C. Franken, A. Geluk, K. E. van Meijgaarden, S. A. Joosten, W. H. Boom, B. Thiel, H. Mayanja-Kizza, M. Joloba, S. Zalwango, M. Nsereko, B. Okwera, H. Kisingo, S. H. E. Kaufmann, S. K. Parida, R. Golinski, J. Maertzdorf, J. Weiner, M. Jacobson, H. Dockrell, S. Smith, P. Gorak-Stolinska, Y.-G. Hur, M. Lalor, J.-S. Lee, A. C. Crampin, N. French, B. Ngwira, A. Ben-Smith, K. Watkins, L. Ambrose, F. Simukonda, H. Mvula, F. Chilongo, J. Saul, K. Branson, S. Suliman, T. J. Scriba, H. Mahomed, E. J. Hughes, N. Bilek, M. Erasmus, O. Xasa, A. Veldsman, K. Downing, M. Fisher, A. Penn-Nicholson, H. Mulenga, B. Abel, M. Bowmaker, B. Kagina, W. K. Chung, W. A. Hanekom, J. Sadoff, D. Sizemore, S. Ramachandran, L. Barker, M. Brennan, F. Weichold, S. Muller, L. Geiter, D. Kassa, A. Abebe, T. Mesele, B. Tegbaru, D. van Baarle, F. Miedema, R. Howe, A. Mihret, A. Aseffa, Y. Bekele, R. Iwnetu, M. Tafesse, L. Yamuah, M. Ota, J. Sutherland, P. Hill, R. Adegbola, T. Corrah, M. Antonio, T. Togun, I. Adetifa, S. Donkor, P. Andersen, I. Rosenkrands, M. Doherty, K. Weldingh, G. Schoolnik, G. Dolganov, T. Van, F. Kafaar, L. Workman, H. Mulenga, T. J. Scriba, E. J. Hughes, N. Bilek, M. Erasmus, O. Xasa, A. Veldsman, Y. Cloete, D. Abrahams, S. Moyo, S. Gelderbloem, M. Tameris, H. Geldenhuys, W. Hanekom, G. Hussey, R. Ehrlich, S. Verver, L. Geiter, Four-Gene Pan-African Blood Signature Predicts Progression to Tuberculosis, Am. J. Respir. Crit. Care Med. 197, 1198–1208 (2018).

23. World Health Organization, Development of a Target Product Profile (TPP) and a framework for evaluation for a test for predicting progression from tuberculosis infection to active disease (World Health Organization, Geneva, Switzerland; http://apps.who.int/iris/bitstream/handle/10665/259176/WHO-HTM-TB-2017.18-eng.pdf;jsessionid=EBD2B5F9B500750ECB57D8E796BFD533?sequence=1), p. (WHO, Geneva, 2017).

24. H. P. Gideon, J. A. Skinner, N. Baldwin, J. L. Flynn, P. L. Lin, Early Whole Blood Transcriptional Signatures Are Associated with Severity of Lung Inflammation in Cynomolgus Macaques with Mycobacterium tuberculosis Infection, J. Immunol. 197, 4817–4828 (2016).

25. R. R. Chowdhury, F. Vallania, Q. Yang, C. J. L. Angel, F. Darboe, A. Penn-Nicholson, V. Rozot, E. Nemes, S. T. Malherbe, K. Ronacher, G. Walzl, W. Hanekom, M. M. Davis, J. Winter, X. Chen, T. J. Scriba, P. Khatri, Y. Chien, A multi-cohort study of the immune factors associated with M. tuberculosis infection outcomes, Nature 560, 644–648 (2018).

26. M. Gonzalez-Juarrero, L. C. Kingry, D. J. Ordway, M. Henao-Tamayo, M. Harton, R. J. Basaraba, W. H. Hanneman, I. M. Orme, R. A. Slayden, Immune Response to Mycobacterium tuberculosis and Identification of Molecular Markers of Disease, Am. J. Respir. Cell Mol. Biol. 40, 398–409 (2009).

27. H.-J. Mollenkopf, K. Hahnke, S. H. E. Kaufmann, Transcriptional responses in mouse lungs induced by vaccination with Mycobacterium bovis BCG and infection with Mycobacterium tuberculosis, Microbes Infect. 8, 136–144 (2006).

28. L. Shi, H. Salamon, E. A. Eugenin, R. Pine, A. Cooper, M. L. Gennaro, Infection with *Mycobacterium tuberculosis* induces the Warburg effect in mouse lungs, Sci. Rep. 5, srep18176 (2015).

29. G. L. Beamer, J. Turner, Murine models of susceptibility to tuberculosis, Arch. Immunol. Ther. Exp. (Warsz.) 53, 469–483 (2005).

30. Medina, North, Resistance ranking of some common inbred mouse strains to Mycobacterium tuberculosis and relationship to major histocompatibility complex haplotype and Nramp1 genotype, Immunology 93, 270–274 (1998).

31. W. P. Gill, N. S. Harik, M. R. Whiddon, R. P. Liao, J. E. Mittler, D. R. Sherman, A replication clock for *Mycobacterium tuberculosis*, Nat. Med. 15, 211–214 (2009).

32. S. V. Capuano, D. A. Croix, S. Pawar, A. Zinovik, A. Myers, P. L. Lin, S. Bissel, C. Fuhrman, E. Klein, J. L. Flynn, Experimental Mycobacterium tuberculosis Infection of Cynomolgus Macaques Closely Resembles the Various Manifestations of Human M. tuberculosis Infection, Infect. Immun. 71, 5831–5844 (2003).

33. P. L. Lin, M. Rodgers, L. Smith, M. Bigbee, A. Myers, C. Bigbee, I. Chiosea, S. V. Capuano, C. Fuhrman, E. Klein, J. L. Flynn, Quantitative Comparison of Active and Latent Tuberculosis in the Cynomolgus Macaque Model, Infect. Immun. 77, 4631–4642 (2009).

34. A. Poulsen, Some clinical features of tuberculosis, Acta Tuberc. Scand. 33, 37–92; concl (1957).

35. T. Gedde-Dahl, Tuberculous infection in the light of tuberculin matriculation, Am. J. Hyg. 56, 139–214 (1952).

36. O. R. McCarthy, Asian immigrant tuberculosis--the effect of visiting Asia, Br. J. Dis. Chest 78, 248–253 (1984).

37. H.-A. Hatherell, X. Didelot, S. L. Pollock, P. Tang, A. Crisan, J. C. Johnston, C. Colijn, J. L. Gardy, Declaring a tuberculosis outbreak over with genomic epidemiology, Microb. Genomics 2 (2016), doi:10.1099/mgen.0.000060.

38. K. Dijkman, C. C. Sombroek, R. A. W. Vervenne, S. O. Hofman, C. Boot, E. J. Remarque, C. H. M. Kocken, T. H. M. Ottenhoff, I. Kondova, M. A. Khayum, K. G. Haanstra, M. P. M. Vierboom, F. A. W. Verreck, Prevention of tuberculosis infection and disease by local BCG in repeatedly exposed rhesus macaques, Nat. Med. 25, 255 (2019).

39. J. R. Andrews, F. Noubary, R. P. Walensky, R. Cerda, E. Losina, C. R. Horsburgh, Risk of Progression to Active Tuberculosis Following Reinfection With Mycobacterium tuberculosis, Clin. Infect. Dis. 54, 784–791 (2012).

40. F. Vallania, A. Tam, S. Lofgren, S. Schaffert, T. D. Azad, E. Bongen, W. Haynes, M. Alsup, M. Alonso, M. Davis, E. Engleman, P. Khatri, Leveraging heterogeneity across multiple datasets increases cell-mixture deconvolution accuracy and reduces biological and technical biases, Nat. Commun. 9, 4735 (2018).

41. T. J. Scriba, A. Penn-Nicholson, S. Shankar, T. Hraha, E. G. Thompson, D. Sterling, E. Nemes, F. Darboe, S. Suliman, L. M. Amon, H. Mahomed, M. Erasmus, W. Whatney, J. L. Johnson, W. H. Boom, M. Hatherill, J. Valvo, M. A. D. Groote, U. A. Ochsner, A. Aderem, W. A. Hanekom, D. E. Zak, other members of the A. cohort study Team, Sequential inflammatory processes define human progression from M. tuberculosis infection to tuberculosis disease, PLOS Pathog. 13, e1006687 (2017).

42. T. Domaszewska, L. Scheuermann, K. Hahnke, H. Mollenkopf, A. Dorhoi, S. H. E. Kaufmann, J. Weiner, Concordant and discordant gene expression patterns in mouse strains identify best-fit animal model for human tuberculosis, Sci. Rep. 7, 1–13 (2017).

43. W. A. Haynes, F. Vallania, C. Liu, E. Bongen, A. Tomczak, M. Andres-Terrè, S. Lofgren, A. Tam, C. A. Deisseroth, M. D. Li, T. E. Sweeney, P. Khatri, Empowering Multi-Cohort Gene Expression Analysis to Increase Reproducibility, Pac. Symp. Biocomput. Pac. Symp. Biocomput. 22, 144–153 (2016).

44. A. Poulsen, Some clinical features of tuberculosis. 1. Incubation period, Acta Tuberc. Scand. 24, 311–346 (1950).

45. F. Cobelens, S. Kik, H. Esmail, D. M. Cirillo, C. Lienhardt, A. Matteelli, From latent to patent: rethinking prediction of tuberculosis, Lancet Respir. Med. 5, 243–244 (2017).

46. R. K. Gupta, C. T. Turner, C. Venturini, H. Esmail, M. X. Rangaka, A. Copas, M. Lipman, I. Abubakar, M. Noursadeghi, Concise whole blood transcriptional signatures for incipient tuberculosis: A systematic review and patient-level pooled meta-analysis, bioRxiv, 668137 (2019).

47. Y. Luo, S. Suliman, S. Asgari, T. Amariuta, Y. Baglaenko, M. Martínez-Bonet, K. Ishigaki, M. Gutierrez-Arcelus, R. Calderon, L. Lecca, S. R. León, J. Jimenez, R. Yataco, C. Contreras, J. T. Galea, M. Becerra, S. Nejentsev, P. A. Nigrovic, D. B. Moody, M. B. Murray, S. Raychaudhuri, Early progression to active tuberculosis is a highly heritable trait driven by 3q23 in Peruvians, Nat. Commun. 10, 1–10 (2019).

48. N. du Plessis, L. Loebenberg, M. Kriel, F. von Groote-Bidlingmaier, E. Ribechini, A. G. Loxton, P. D. van Helden, M. B. Lutz, G. Walzl, Increased Frequency of Myeloid-derived Suppressor Cells during Active Tuberculosis and after Recent Mycobacterium tuberculosis Infection Suppresses T-Cell Function, Am. J. Respir. Crit. Care Med. 188, 724–732 (2013).

49. J. Weiner, T. Domaszewska, S. Donkor, S. H. E. Kaufmann, P. C. Hill, J. S. Sutherland, Changes in transcript, metabolite and antibody reactivity during the early protective immune response in humans to Mycobacterium tuberculosis infection, Clin. Infect. Dis., doi:10.1093/cid/ciz785.

50. R. Chen, L. Xia, K. Tu, M. Duan, K. Kukurba, J. Li-Pook-Than, D. Xie, M. Snyder, Longitudinal personal DNA methylome dynamics in a human with a chronic condition, Nat. Med. 24, 1930–1939 (2018).

51. H. Esmail, R. Lai, M. Lesosky, K. Wilkinson, C. Graham, A. Coussens, T. Oni, J. Warwick, Q. Said-Hartley, C. Koegelenberg, G. Walzl, J. Flynn, D. Young, C. Barry, A. O’Garra, R. Wilkinson, Characterization of progressive HIV-associated tuberculosis using 2-deoxy-2-[18F]fluoro-D-glucose positron emission and computed tomography, Nat. Med. 22, 1090–1093 (2016).

52. E. J. Hwang, S. Park, K.-N. Jin, J. I. Kim, S. Y. Choi, J. H. Lee, J. M. Goo, J. Aum, J.-J. Yim, C. M. Park, D. H. Kim, W. Woo, C. Choi, I. P. Hwang, Y. S. Song, L. Lim, K. Kim, J. Y. Wi, S. S. Oh, M.-J. Kang, Development and Validation of a Deep Learning–based Automatic Detection Algorithm for Active Pulmonary Tuberculosis on Chest Radiographs, Clin. Infect. Dis. 69, 739–747 (2019).

53. M. T. Coleman, P. Maiello, J. Tomko, L. J. Frye, D. Fillmore, C. Janssen, E. Klein, P. L. Lin, Early Changes by 18Fluorodeoxyglucose Positron Emission Tomography Coregistered with Computed Tomography Predict Outcome after Mycobacterium tuberculosis Infection in Cynomolgus Macaques, Infect. Immun. 82, 2400–2404 (2014).

54. P. L. Lin, J. L. Flynn, The End of the Binary Era: Revisiting the Spectrum of Tuberculosis, J. Immunol. 201, 2541–2548 (2018).

55. A. Singhania, R. J. Wilkinson, M. Rodrigue, P. Haldar, A. O’Garra, The value of transcriptomics in advancing knowledge of the immune response and diagnosis in tuberculosis, Nat. Immunol. 19, 1159–1168 (2018).

56. O. Van Der Meeren, M. Hatherill, V. Nduba, R. J. Wilkinson, M. Muyoyeta, E. Van Brakel, H. M. Ayles, G. Henostroza, F. Thienemann, T. J. Scriba, A. Diacon, G. L. Blatner, M.-A. Demoitié, M. Tameris, M. Malahleha, J. C. Innes, E. Hellström, N. Martinson, T. Singh, E. J. Akite, A. Khatoon Azam, A. Bollaerts, A. M. Ginsberg, T. G. Evans, P. Gillard, D. R. Tait, Phase 2b Controlled Trial of M72/AS01E Vaccine to Prevent Tuberculosis, N. Engl. J. Med. 379, 1621–1634 (2018).

57. L. L. Lu, M. T. Smith, K. K. Q. Yu, C. Luedemann, T. J. Suscovich, P. S. Grace, A. Cain, W. H. Yu, T. R. McKitrick, D. Lauffenburger, R. D. Cummings, H. Mayanja-Kizza, T. R. Hawn, W. H. Boom, C. M. Stein, S. M. Fortune, C. Seshadri, G. Alter, IFN-γ-independent immune markers of Mycobacterium tuberculosis exposure, Nat. Med., 1 (2019).

58. J. Weiner, J. Maertzdorf, J. S. Sutherland, F. J. Duffy, E. Thompson, S. Suliman, G. McEwen, B. Thiel, S. K. Parida, J. Zyla, W. A. Hanekom, R. P. Mohney, W. H. Boom, H. Mayanja-Kizza, R. Howe, H. M. Dockrell, T. H. M. Ottenhoff, T. J. Scriba, D. E. Zak, G. Walzl, S. H. E. Kaufmann, Metabolite changes in blood predict the onset of tuberculosis, Nat. Commun. 9, 5208 (2018).

59. B. Vesosky, E. K. Rottinghaus, C. Davis, J. Turner, CD8 T Cells in Old Mice Contribute to the Innate Immune Response to Mycobacterium tuberculosis via Interleukin-12p70-Dependent and Antigen-Independent Production of Gamma Interferon, Infect. Immun. 77, 3355–3363 (2009).

60. J. C. Cyktor, B. Carruthers, P. Stromberg, E. Flaño, H. Pircher, J. Turner, Killer Cell Lectin-Like Receptor G1 Deficiency Significantly Enhances Survival after Mycobacterium tuberculosis Infection, Infect. Immun. 81, 1090–1099 (2013).

61. A. Lachmann, D. Torre, A. B. Keenan, K. M. Jagodnik, H. J. Lee, L. Wang, M. C. Silverstein, A. Ma’ayan, Massive mining of publicly available RNA-seq data from human and mouse, Nat. Commun. 9, 1366 (2018).

62. A. Liaw, M. Wiener, Classification and regression by random forest, R News 2, 18–22 (2002).

63. M. Kuhn, J. Wing, S. Weston, A. Williams, C. Keefer, A. Engelhardt, T. Cooper, Z. Mayer, B. Kenkel, the R. C. Team, M. Benesty, R. Lescarbeau, A. Ziem, L. Scrucca, Y. Tang, C. Candan, and T. Hunt, caret: Classification and Regression Training (https://CRAN.R-project.org/package=caret, 2018; https://CRAN.R-project.org/package=caret).

64. M. N. Wright, A. Ziegler, ranger: A Fast Implementation of Random Forests for High Dimensional Data in C++ and R, J. Stat. Softw. 77 (2017), doi:10.18637/jss.v077.i01.

65. B. Greenwell, B. Boehmke, J. Cunningham, G. B. M. Developers (https://github.com/gbm-developers), *gbm: Generalized Boosted Regression Models* (https://CRAN.R-project.org/package=gbm, 2018; https://CRAN.R-project.org/package=gbm).

66. A. Karatzoglou, A. Smola, K. Hornik, A. Zeileis, kernlab - An S4 Package for Kernel Methods in R, J. Stat. Softw. 11, 1–20 (2004).

67. J. Friedman, T. Hastie, R. Tibshirani, Regularization Paths for Generalized Linear Models via Coordinate Descent, J. Stat. Softw. 33, 1–22 (2010).

68. T. E. Sweeney, L. Braviak, C. M. Tato, P. Khatri, Genome-wide expression for diagnosis of pulmonary tuberculosis: a multicohort analysis, Lancet Respir. Med. 4, 213–224 (2016).

69. S. Durinck, P. T. Spellman, E. Birney, W. Huber, Mapping identifiers for the integration of genomic datasets with the R/Bioconductor package biomaRt, Nat. Protoc. 4, 1184–1191 (2009).

70. S. Li, N. Rouphael, S. Duraisingham, S. Romero-Steiner, S. Presnell, C. Davis, D. S. Schmidt, S. E. Johnson, A. Milton, G. Rajam, S. Kasturi, G. M. Carlone, C. Quinn, D. Chaussabel, A. K. Palucka, M. J. Mulligan, R. Ahmed, D. S. Stephens, H. I. Nakaya, B. Pulendran, Molecular signatures of antibody responses derived from a systems biology study of five human vaccines, Nat. Immunol. 15, 195–204 (2014).

71. T. Domaszewska, J. Weiner, disco: Discordance and Concordance of Transcriptomic Responses (2018; https://CRAN.R-project.org/package=disco).

72. J. Weiner 3rd, T. Domaszewska, tmod: an R package for general and multivariate enrichment analysis (PeerJ Preprints, 2016; https://peerj.com/preprints/2420).

73. 73. NCAR, verification: Weather Forecast Verification Utilities (https://CRAN.R-project.org/package=verification, 2015; https://CRAN.R-project.org/package=verification).

74. X. Robin, N. Turck, A. Hainard, N. Tiberti, F. Lisacek, J.-C. Sanchez, M. Müller, pROC: an open-source package for R and S+ to analyze and compare ROC curves, BMC Bioinformatics 12, 77–84 (2011).

75. D. Bates, M. Mächler, B. Bolker, S. Walker, Fitting Linear Mixed-Effects Models Using lme4, J. Stat. Softw. 67, 1–48 (2015).

